# Characterization of nick binding and sealing by LIG1 Huntington’s disease-asssociated K845N variant at biochemical, structural, and single-molecule levels

**DOI:** 10.1101/2025.04.26.650789

**Authors:** Jacob E. Ratcliffe, Kanal E. Balu, Surajit Chatterjee, Kar Men Lee, Melike Çağlayan

## Abstract

DNA ligase 1 (LIG1) joins broken strand breaks and discriminates against nicks containing mismatch or oxidative damage. Huntington’s disease (HD)-associated mutation K845N in *LIG1* gene has been predicted to be onset delaying and suppresses CAG repeat expansion. Yet, how this mutation impacts faithful nick sealing and efficient DNA binding by LIG1 remains unknown. Here, using biochemistry, X-ray crystallography, and total internal reflection fluorescence microscopy, we comprehensively characterized the impact of LIG1 HD-associated mutation at biochemical, structural, and single-molecule levels. Our results showed a reduced ligation efficiency by LIG1 K845N variant in the presence of nick substrates containing all possible 12 mismatches, 8-oxoG, and ribonucleotides at the 3’-end when compared with the wild-type enzyme. Furthermore, our structures provided an atomic insight into differences in distances between the functional groups of K/N845 and DNA ends, demonstrating similar conformation and a lack of large scale alternations at the ligase active site. Finally, our single-molecule measurements in real-time revealed that K845N mutant binds less frequently for shorter life-time to nick DNA than wild-type protein. Overall findings contribute to understand the mechanism by which LIG1 ensures fidelity and nick binding to maintain genome stability at the final ligation step in normal *versus* disease states.

## Introduction

Mammalian genomes can be exposed to multiple types of DNA damage that can be generated during endogenous cellular processes or by the effect of exogenous sources such as environmental factors including UV-light, chemical toxins, irradiation, inflammation, and nutritional factors (1). The combined actions of DNA replication machinery and a network of DNA repair pathways are essential for maintaining overall genome stability and DNA ligases seal nicks to repair broken strand breaks during these criticial DNA transactions (2). ATP-dependent human DNA ligases catalyze a phosphodiester bond formation between the 3’-hydroxyl (OH) and 5’-phosphate (PO_4_) ends of nick in a conserved a three-step of DNA ligation reaction (3). In the first step, the formation of the covalent ligase-adenylate intermediate occurs after DNA ligase reacts with a nucleotide cofactor and releases a pyrophosphate group, which is followed by transfer of an adenosine monophosphate (AMP) moiety is from the adenylated ligase to the 5’-PO_4_ end of nick in the second step (4). In the final step of the ligation reaction, using 3’-OH as a nucleophile, and the ligase seals DNA ends which is coupled to a release of an AMP (5).

Of the three human DNA ligases, DNA ligase 1 (LIG1) is the main replicative ligase playing a critical role for the maturation of Okazaki fragments with >50 million ligation events during DNA replication and is responsible for the majority of DNA ligase activity in proliferating cells (6, 7). Furthermore, LIG1 finalizes DNA excision repair pathways with an ultimate nick sealing step (8). The expression level of LIG1 was found higher in human cancer cells and the ligase is also essential for embryonic development while some studies have also demonstrated that LIG1 is not required for cellular DNA replication and that either LIG3 or LIG4 can substitute for joining Okazaki fragments (9–11).

The fidelity of ligation at the final step of DNA repair pathway requires Watson-Crick base-paired end at the nick (*i.e*. 3’-dG:C) after an incorporation of a correct nucleotide in a gap repair intermediate by DNA polymerase (*i.e*., dGTP:C) during DNA synthesis step (12). However, in the presence of a modified base at the 3’-end, LIG1 fails resulting in the formation of ligase failure or abortive ligation products with a 5’-adenylate (AMP) group (13). In our previous studies, we demonstrated that, after incorporation of mismatches or oxidized nucleotides (8-oxodGTP) by DNA polymerase (pol) β at the downstream steps of Base Excision Repair (BER) pathway, resulting nick product with oxidatively damaged base, 8-oxo-2’-deoxyguanosine (8-oxodG), or all 12 possible non-canonical mismatches at the 3’-end cannot be efficiently sealed by LIG1 (14–22). Our LIG1 structures also demonstrated the strategies that the ligase active site utilizes to discriminate against nicks with mismatches depending on the architecture of 3’-terminus:template base-pairing (23). Similarly, our structures revealed the mechanism by which LIG1 handles nicks with oxidatively damaged end by undergoing distinct adjustments around nucleotides relative to the nick site depending on the dual coding potential of 8oxoG in -*anti* versus -*syn* conformation (24). Furthermore, in our structures of LIG1/RNA-DNA hybrids, we previously demonstrated that the ligase shows a lack of discrimination against nicks containing a single ribonucleotide at the 3’-end and can seal almost all possible 3’-ribonucleotide-containing mismatches as efficient as nicks with Watson-Crick base-paired ends (25, 26). These studies contribute to understand the mechanism by which LIG1 ensures fidelity while engaging with potentially mutagenic and toxic repair intermediates at the final step. However, it’s still largely unknown how pertubrations at the ligase active site, particularly disease-associated mutations in the catalytic core of LIG1, could impact the accuracy of nick sealing and efficiency of DNA binding.

The reiteration of mutations identified in the *LIG1* gene of human individuals, P529L, E566K, R641L, and R771W, whose symptoms included developmental abnormalities, immunodeficiency and lymphoma, in a mouse model has proven that LIG1 deficiency causes genomic instability and predisposition to cancer (27–29). Furthermore, LIG1^-/-^ HEK293T cells cells complemented with LIG1 syndrome alleles show hypersensitivity to DNA damaging agents and a deficiency in DNA repair (30–32). Polyglutamine disorders are inheritable and are caused by the expansion of Cytosine-Adenine-Guanine (CAG) repeats in DNA (33). Huntington’s disease (HD) is the most frequent of the polyglutamine diseases as a dominantly inherited neurological disorder, characterized by an array of symptoms such as cognitive decline, involuntary movements, and changes to behavior and speech (34). The expansion of CAG repeats in the Huntingtin (HTT) gene leads to toxic Huntingtin protein (mHTT) and its build up disrupts function of neurons and neurodegeneration (35). HD is caused by unstable repeat expansion and lengthening of the trinucleotide repeat CAG (36). The Genetic Modifiers of HD (GeM-HD) Consortium utilized a genome wide association study (GWAS) of over 9, 000 individuals with HD and reported that HD age at onset is determined by CAG repeat length regardless of the length of polyglutamine, demonstrating a relationship between CAG repeats and HD symptom onset (37). It has been reported in this study that the chromosome 19 that encodes LIG1 reveals a rare modifier haplotype (19AM3) that is strongly onset delaying and is tagged by LIG1 missense change K845N, suggesting that reduced activity and/or altered interactions of LIG1 protein may suppress CAG repeat expansion (37). Yet, it remains unknown how the HD-associated K845N mutation in *LIG1* gene could impact nick binding and sealing efficiency in the presence of non-canonical substrates that mimic base-substitution errors or potentially mutagenic repair intermediates introduced by repair and replication DNA polymerases, which is critical to elucidate how LIG1 ensures fidelity in normal *versus* disease states.

In the present study, using a combined approach including biochemistry, X-ray crystallography, and total internal reflection fluorescence (TIRF) microscopy, we explored the impact of LIG1 HD-associated mutation K845N on the efficiency of nick sealing and binding at biochemical, structural, and single-molecule levels. For this purpose, we tested end joining ability of LIG1 wild-type and K845N variant for the nick DNA substrates containing canonical, mismatched, damaged, and ribonucleotide-containing ends *in vitro*. Our findings demonstrated that K845N mutation impacts the ligation efficiency of wild-type enzyme for a variety of nick DNA substrates and we mostly obtained reduced amount of ligation products. Furthermore, to understand the structural arrangements at the ligase/nick DNA interactions by the effect of HD-associated mutation, we determined X-ray structure of LIG1 K845N mutant in complex with nick DNA containing canonical end. Our structures showed a similarity in the DNA conformation at the downstream of the nick and distances relative to the 3’-and 5’-ends in the absence and presence of K845N mutation. Finally, our single-molecule nick DNA binding measurements in real-time demonstrated that K845N mutant binds less frequently for shorter time period to nick when compared to LIG1 wild-type. Overall, our findings provide an insights into the mechanism by LIG1 HD-associated K845N variant impacts the accuracy of final ligation step. This study could contribute to elucidate the biochemical mechanism of the neurodegenerative symptoms caused by HD and biochemical defects observed in LIG1 pathological variant underscore LIG1 function in distinct disease states.

## Results

### Impact of LIG1 K845N mutation on the ligation efficiency of nick DNA substrates containing canonical, mismatched and damaged ends

We first comprehensively characterized the ligation profile of LIG1 K845N variant for the nick DNA substrates containing all possible 12 non-canonical mismatches at the 3’-end. These substrates are biologically relevant, and mimic base substitution errors that can be inserted by DNA polymerases during repair and replication (Supplementary Scheme 1A). Consistent with our previous report (15, 23), we showed that LIG1 wild-type can seal the nick DNA substrates containing mismatches at different efficiency depending on the architecture of the 3’-terminus:template base pairing (Supplementary Figures 1-2). Similarly, we observed subtle changes in the ligation efficiency of LIG1 K845N variant that exhibits relatively less efficient nick sealing for almost all mismatches (Figures 1-2). For template A mismatches, we observed higher ligation efficiency for 3’-dC:A mismatch, while both LIG1 wild-type and K845N show a similar ligation profile for nick DNA substrate with 3’-dA:A mismatch (Figure 1A, C and Supplementary Figure 1A, C). However, nick DNA containing purine:purine mismatch 3’-dG:A cannot be ligated by either LIG1 protein. In case of nick DNA substrates containing template T mismatches, we observed significantly reduced ligation efficiency in the presence of 3’-dG:T mismatch, while end joining of the nick with 3’-dC:T mismatch was relatively lower by LIG1 K845N (Figure 1B, D). Interestingly, nick DNA substrate with 3’-dT:T mismatch cannot be sealed by either LIG1 protein (Supplementary Figure 1B, D). Regarding nick DNA substrates containing template G mismatches 3’-dG:G, 3’-dA:G, and 3’-dT:G, we overall obtained inefficient ligation by LIG1 wild-type and K845N mutant (Figure 2A, C and Supplementary Figure 2A, C). For template C mismatches, both LIG1 proteins have around 80% of ligation product in the presence of nick substrate with 3’-dT:C mismatch (Figure 2B, D and Supplementary Figure 2B, D). However, in the presence of 3’-dC:C and 3’-dA:C mismatches, we obtained an inefficient ligation by LIG1 wild-type, which was further reduced by the effect of K845N mutation. For all possible 12 mismatches, we obtained the highest amount of DNA-AMP intermediate formation in the presence of nick substrates with 3’-dG:G mismatch by LIG1 K845N, suggesting an inability to incorporate bulky purine-purine bases resulting in the formation of more ligation failure products with a 5’-AMP (Supplementary Figures 3-4). We obserbed the ligation of all four canonical nick substrates by LIG1 K845N mutant (Supplementary Figures 5-6).

**Figure 1.**
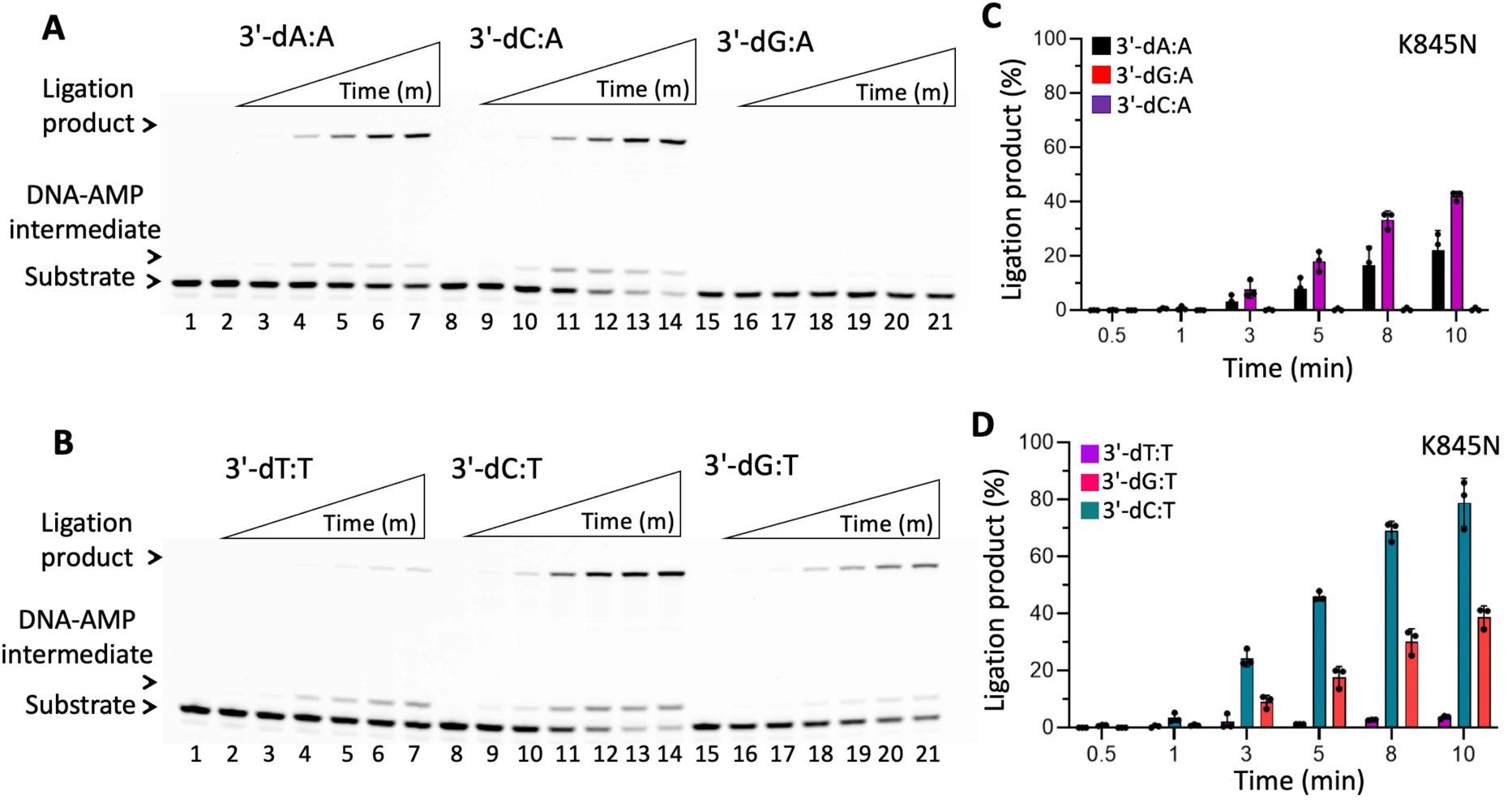
Ligation efficiency of LIG1 K845N mutant for nick DNA substrates containing template A and T mismatches. **(A)** Lanes 1, 8, and 15 are the negative enzyme controls of the nick DNA substrates with 3’-dA:A, 3’-dC:A, and 3’-dG:A mismatches, respectively. **(B)** Lanes 1, 8, and 15 are the negative enzyme controls of the nick DNA substrates with 3’-dT:T, 3’-dC:T, and 3’-dG:T mismatches, respectively. In both panels, lanes 2-7, 9-14, and 16-21 are the ligation reaction products by LIG1 K845N mutant, and correspond to time points of 0.5, 1, 3, 5, 8, and 10 min. **(C-D)** Graphs show time-dependent changes in the amount of ligation products. The data represent the average from three independent experiments ± SD.

**Figure 2.**
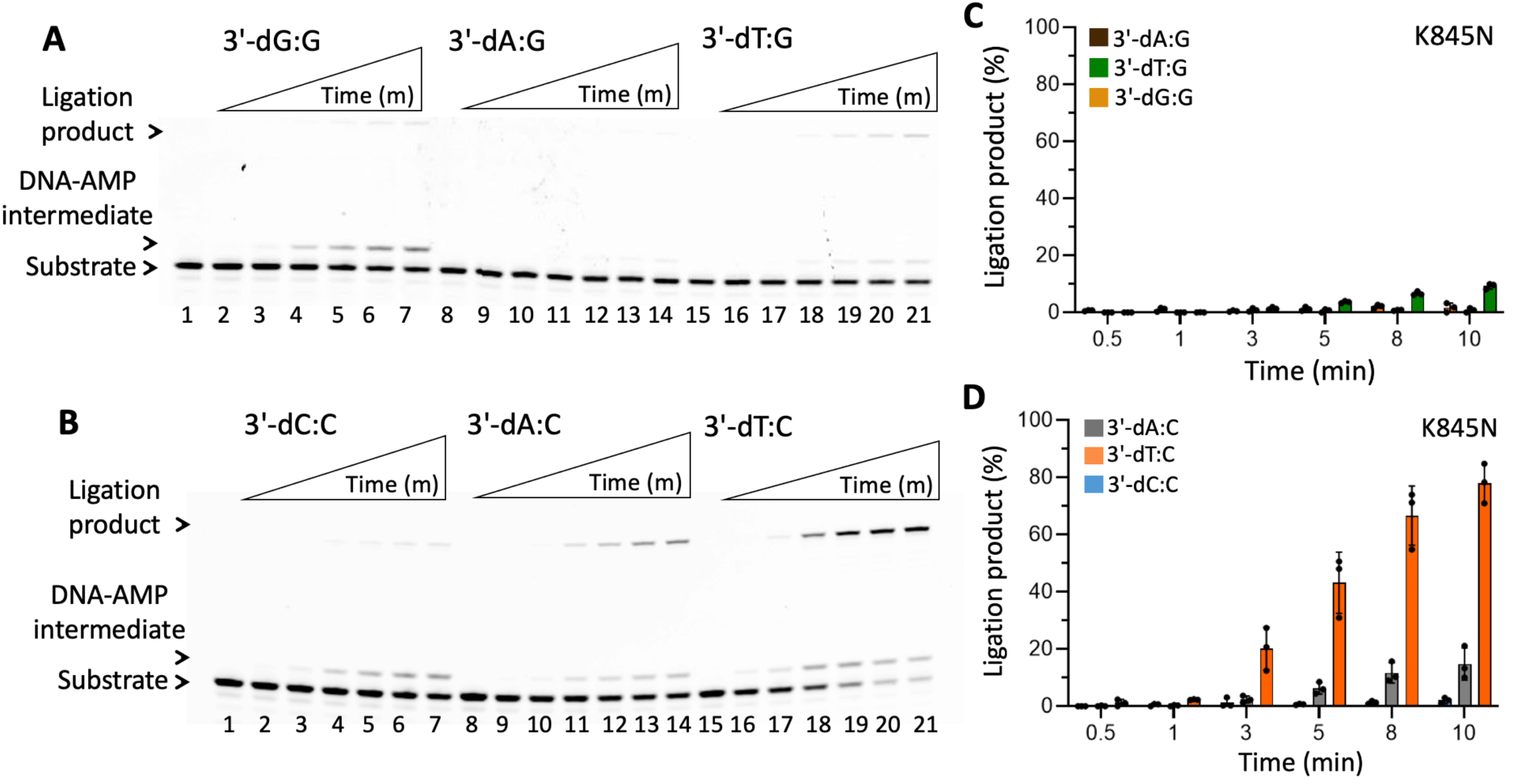
Ligation efficiency of K845N mutant for nick DNA substrates containing template G and C mismatches. **(A)** Lanes 1, 8, and 15 are the negative enzyme controls of the nick DNA substrates with 3’-dG:G, 3’-dA:G, and 3’-dT:G mismatches, respectively. **(B)** Lanes 1, 8, and 15 are the negative enzyme controls of the nick DNA substrates with 3’-dC:C, 3’-dA:C, and 3’-dT:C mismatches, respectively. In both panels, lanes 2-7, 9-14, and 16-21 are the ligation reaction products by LIG1 K845N mutant, and correspond to time points of 0.5, 1, 3, 5, 8, and 10 min. **(C-D)** Graphs show time-dependent changes in the amount of ligation products. The data represent the average from three independent experiments ± SD.

In addition to all 12 mismatches, we characterized the ligation profile of LIG1 K845N mutant for the nick DNA substrates containing damaged ends, particularly 3’-8oxodG opposite template base A or C (Supplementary Scheme 1B). In alignment with our previous reports (15, 24), LIG1 wild-type exhibits the mutagenic ligation of 3’-8oxodG:A and less efficient nick sealing with 3’-8oxodG:C substrate (Supplementary Figure 7A, C). This was also the case for LIG1 K845N mutant (Figure 3A, B), while the mutagenic nick sealing was less efficient at earlier time points of reaction and there was no ligation product in the presence of nick substrate with 3’-8oxodG:C (Figure 3C, D).

**Figure 3.**
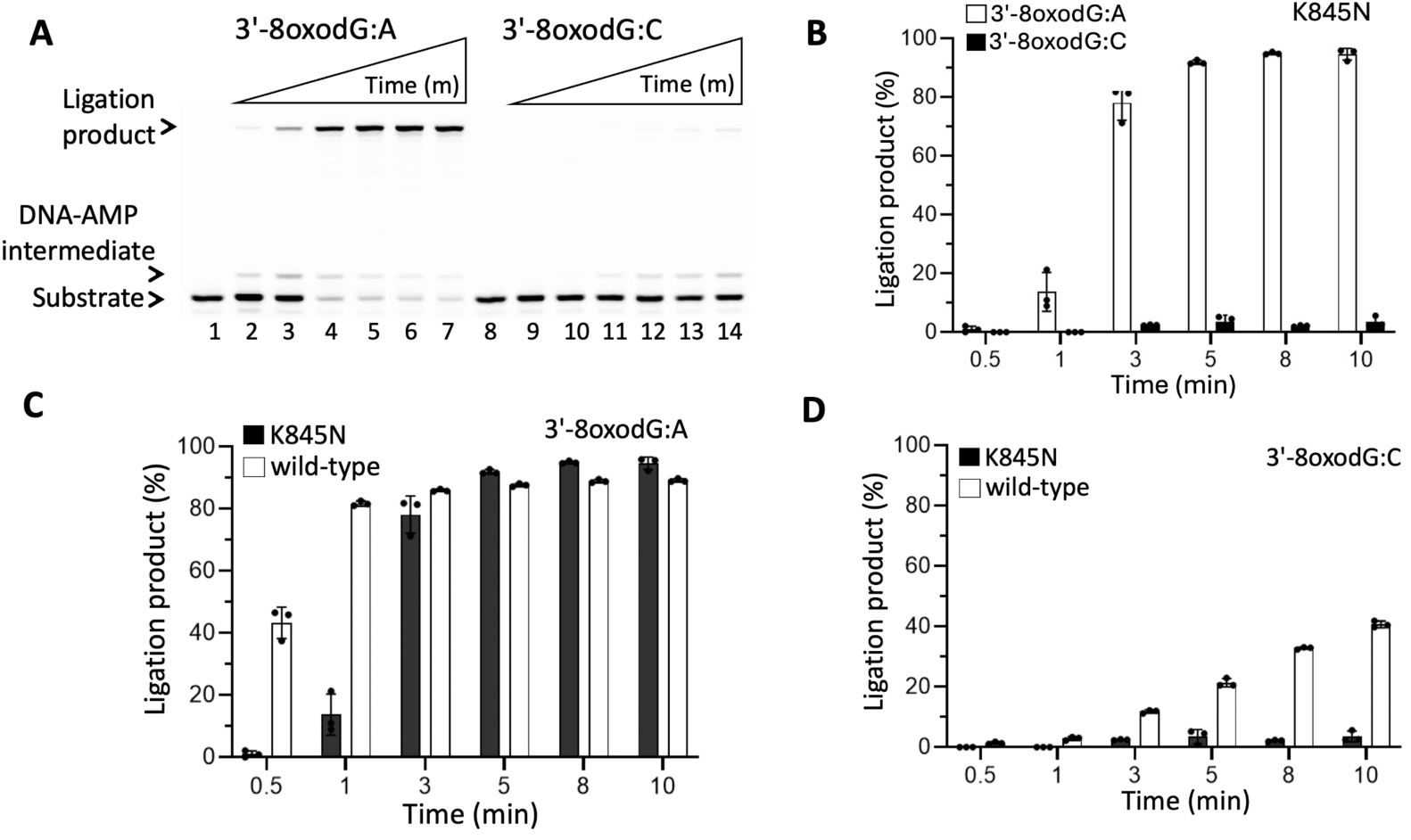
Ligation efficiency of LIG1 K845N mutant for nick DNA substrates containing oxidatively damaged ends. **(A)** Lanes 1 and 8 are the negative enzyme controls of the nick DNA substrates with 3’-8oxodG:A and 3’-8oxodG:C, respectively. Lanes 2-7 and 9-14 are the ligation reaction products by LIG1 K845N mutant in the presence of the nick DNA substrates with 3’-8oxodG:A and 3’-8oxodG:C, respectively, and correspond to time points of 0.5, 1, 3, 5, 8, and 10 min. **(B-D)** Graphs show time-dependent changes in the amount of ligation products by LIG1 K845N mutant and its comparison with wild-type. The data represent the average from three independent experiments ± SD.

### Impact of LIG1 K845N mutation on sugar discrimination against nick DNA containing 3’-ribonucleotides

We also investigated the ligation efficiency of LIG1 K845N variant for the nick DNA substrates containing a single ribonucleotide at the 3’-end (Supplementary Scheme 1C). These nick DNA substrates mimic the ribonucleotide insertion products of DNA polymerases before LIG1 seals the resulting nick intermediate during Okazaki fragment maturation of DNA replication or at the last ligation step of almost all DNA repair pathways (38).

As we previously reported (25, 26), our results with LIG1 wild-type demonstrated a lack of sugar discrimination against a ribonucleotide at the 3’-end of nick DNA. We showed that LIG1 cannot discriminate against a “wrong” sugar and is able to ligate all possible 12 ribonucleotide-containing mismatches (Supplementary Figures 8-9). Similarly, the ligation of nick DNA substrates with 3’-preinserted ribonucleotide mismatches by LIG1 K845N variant yielded efficient ligation (Figures 4-5). For both LIG1 proteins, we observed a robust accumulation of ligation products, especially for the Watson-Crick base paired nick DNA substrates containing 3’-rA:T, 3’-rC:G, and 3’-rG:C. However, there was a relatively less efficient ligation by LIG1 K845N variant for some of the nick DNA substrates containing 3’-ribonucleotides in comparison with LIG1 wild-type such as 3’-rA:C, 3’-rC:C, 3’-rA:G, 3’-rG:T and 3’-rG:A.

**Figure 4.**
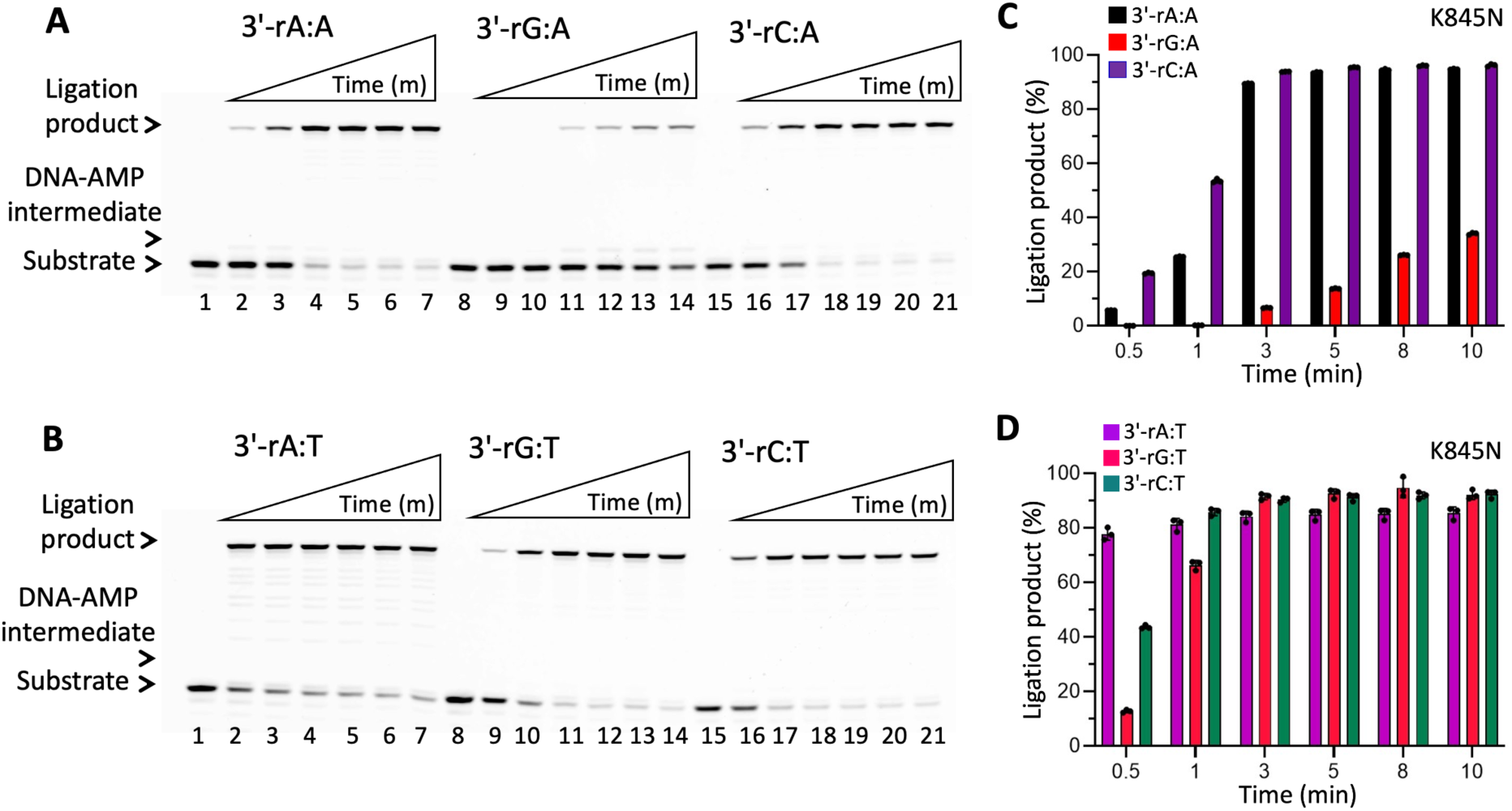
Ligation efficiency of LIG1 K845N mutant for nick DNA substrates containing 3’-ribonucleotides opposite to template A and T. **(A)** Lanes 1, 8, and 15 are the negative enzyme controls of the nick DNA substrates with 3’-rA:A, 3’-rG:A, and 3’-rC:A mismatches, respectively. **(B)** Lanes 1, 8, and 15 are the negative enzyme controls of the nick DNA substrates with 3’-rA:T, 3’-rG:T, and 3’-rC:T mismatches, respectively. In both panels, lanes 2-7, 9-14, and 16-21 are the ligation reaction products by LIG1 K845N mutant, and correspond to time points of 0.5, 1, 3, 5, 8, and 10 min. **(C-D)** Graphs show time-dependent changes in the amount of ligation products. The data represent the average from three independent experiments ± SD.

**Figure 5.**
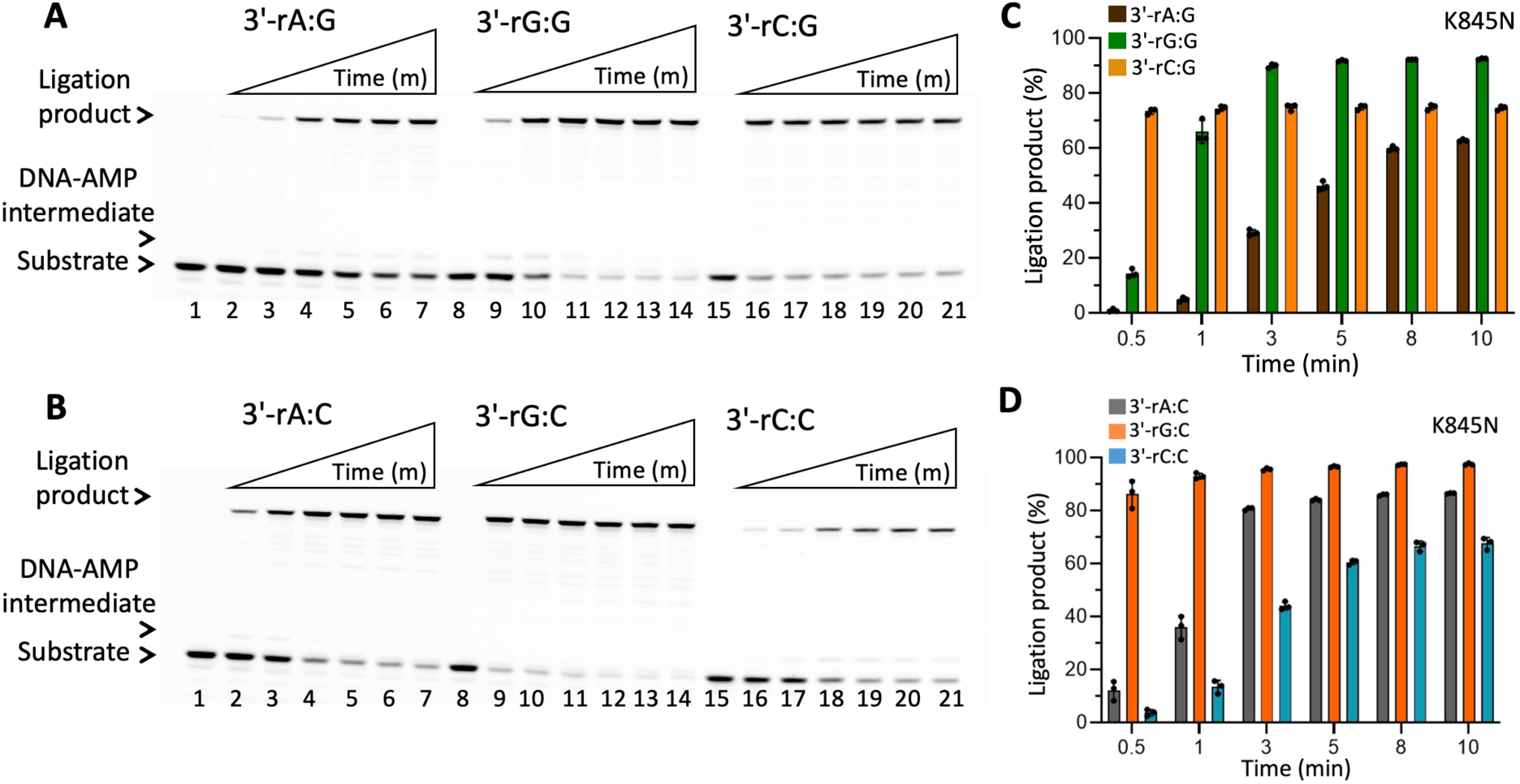
Ligation efficiency of LIG1 K845N mutant for nick DNA substrates containing 3’-ribonucleotides opposite to template G and C. **(A)** Lanes 1, 8, and 15 are the negative enzyme controls of the nick DNA substrates with 3’-rA:G, 3’-rG:G, and 3’-rC:G mismatches, respectively. **(B)** Lanes 1, 8, and 15 are the negative enzyme controls of the nick DNA substrates with 3’-rA:C, 3’-rG:C, and 3’-rC:C mismatches, respectively. In both panels, lanes 2-7, 9-14, and 16-21 are the ligation reaction products by LIG1 K845N mutant, and correspond to time points of 0.5, 1, 3, 5, 8, and 10 min. **(C-D)** Graphs show time-dependent changes in the amount of ligation products. The data represent the average from three independent experiments ± SD.

In addition to all 12 mismatches containing 3’-ribonucleotides, we questioned the ligation efficiency of LIG1 K845N mutant in the presence of the nick DNA substrates containing oxidatively damaged ribonucleotides, particularly 3’-8oxorG opposite template base A or C (Supplementary Scheme 1B). Our results demonstrated the mutagenic ligation of nick substrate with 3’-8oxorG base paired with template A, while the ligation efficiency was relatively less in the presence of nick substrate with 3’-8oxorG:C (Figure 6A, B). The comparison of ligation products between LIG1 wild-type *versus* K845N mutant showed relatively higher amount of nick sealing products by wild-type enzyme (Figure 6C, D and Supplementary Figure 7B, D).

**Figure 6.**
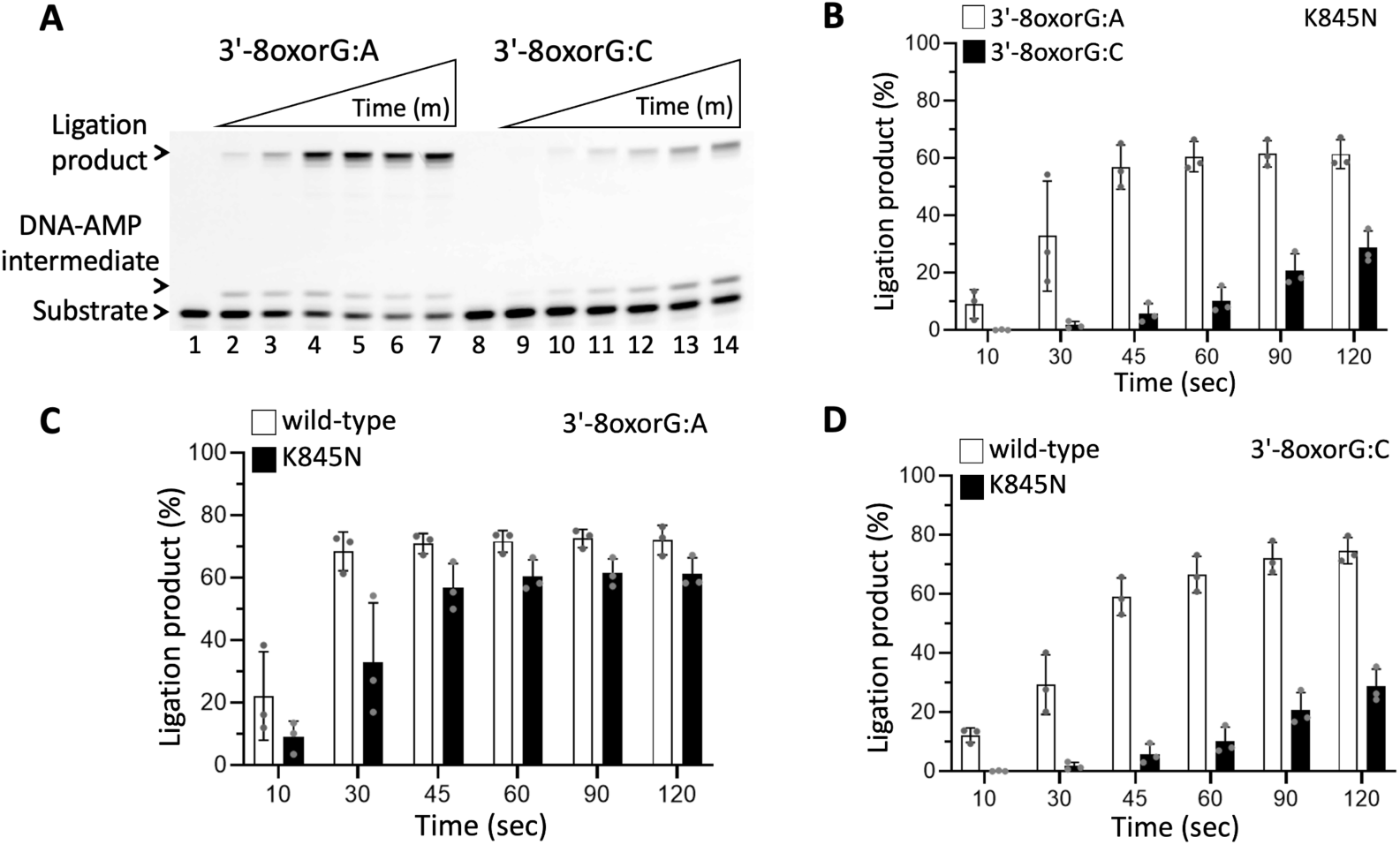
Ligation efficiency of LIG1 K845N mutant for nick DNA substrates containing oxidatively damaged ribonucleotide ends. **(A)** Lanes 1 and 8 are the negative enzyme controls of the nick DNA substrates with 3’-8oxorG:A and 3’-8oxorG:C, respectively. Lanes 2-7 and 9-14 are the ligation reaction products by LIG1 K845N mutant in the presence of the nick DNA substrates with 3’-8oxorG:A and 3’-8oxorG:C, respectively, and correspond to time points of 0.5, 1, 3, 5, 8, and 10 min. **(B-D)** Graphs show time-dependent changes in the amount of ligation products by LIG1 K845N mutant and its comparison with wild-type. The data represent the average from three independent experiments ± SD.

### Structure of LIG1 K845N in complex with nick containing canonical end

In addition to comprehensive biochemical characterization of LIG1 K845N mutant in the ligation assays *in vitro* for the nick DNA substrates containing all 12 non-canonical mismatches, ribonucleotides, and oxidative damage (8oxodG and 8oxorG), we determined the structure of LIG1 K845N in complex with nick DNA containing a canonical A:T (Table 1). For this purpose, we used LIG1 low-fidelity EE/AA mutant that harbors E346A and E592A mutations (LIG1^EE/AA^) for crystallization of the triple mutant LIG1^EE/AA^ K845N (23–26). Also, we used our previously solved structure of LIG1^EE/AA^/A:T for comparison with LIG1^EE/AA^ K845N in the present study (23). There was no significant difference in the ligation efficiency of LIG1^EE/AA^ double-mutant and LIG1^EE/AA^ K845N triple-mutant for sealing of nick substrate with 3’-dA:T (Supplementary Figure 10).

**Table 1:**
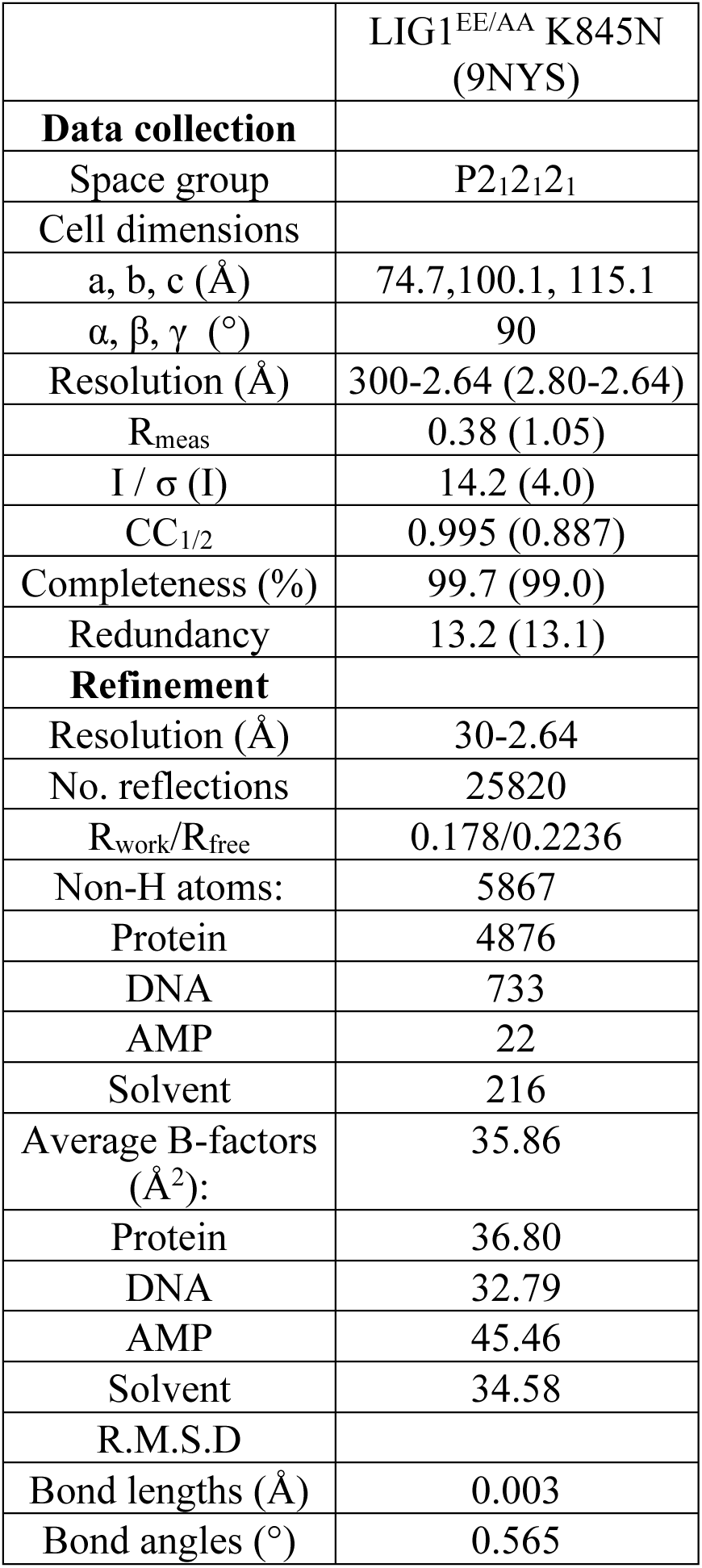
X-ray data collection and refinement statistics of LIG1 structures.

|  |  |
| --- | --- |
|  | LIG1<sup>EE/AA</sup> K845N<br>(9NYS) |
| **Data collection** |  |
| Space group | P2<sub>1</sub>2<sub>1</sub>2<sub>1</sub> |
| Cell dimensions |  |
| a, b, c (Å) | 74.7, 100.1, 115.1 |
| α, β, γ (°) | 90 |
| Resolution (Å) | 300-2.64 (2.80-2.64) |
| R<sub>meas</sub> | 0.38 (1.05) |
| I / σ (I) | 14.2 (4.0) |
| CC<sub>1/2</sub> | 0.995 (0.887) |
| Completeness (%) | 99.7 (99.0) |
| Redundancy | 13.2 (13.1) |
| **Refinement** |  |
| Resolution (Å) | 30-2.64 |
| No. reflections | 25820 |
| R<sub>work</sub>/R<sub>free</sub> | 0.178/0.2236 |
| Non-H atoms: | 5867 |
| Protein | 4876 |
| DNA | 733 |
| AMP | 22 |
| Solvent | 216 |
| Average B-factors<br>(Å<sup>2</sup>): | 35.86 |
| Protein | 36.80 |
| DNA | 32.79 |
| AMP | 45.46 |
| Solvent | 34.58 |
| R.M.S.D |  |
| Bond lengths (Å) | 0.003 |
| Bond angles (°) | 0.565 |

In the structure of LIG1^EE/AA^/A:T, we observed that an AMP moiety is bound to the 5’-end of the nick where the DNA-AMP intermediate is formed, which refers to step 2 of the ligation reaction (Figure 7A). Similarly, in the structure of LIG1^EE/AA^ K845N/A:T, we observed the ligase active site at the step 2 of the ligation reaction (Figure 7B). The overlay of both LIG1 structures demonstrated similar confirmation with the root mean square deviation (RMSD) of 0.560Å. The superimposition of these structures also shows a similarity in the DNA conformation at the downstream of the nick in the absence and presence of K845N mutation residing in the OBD domain (Figure 8). We did not observe any large scale alternations at the ligase active site by the effect of K845N mutation.

**Figure 7.**
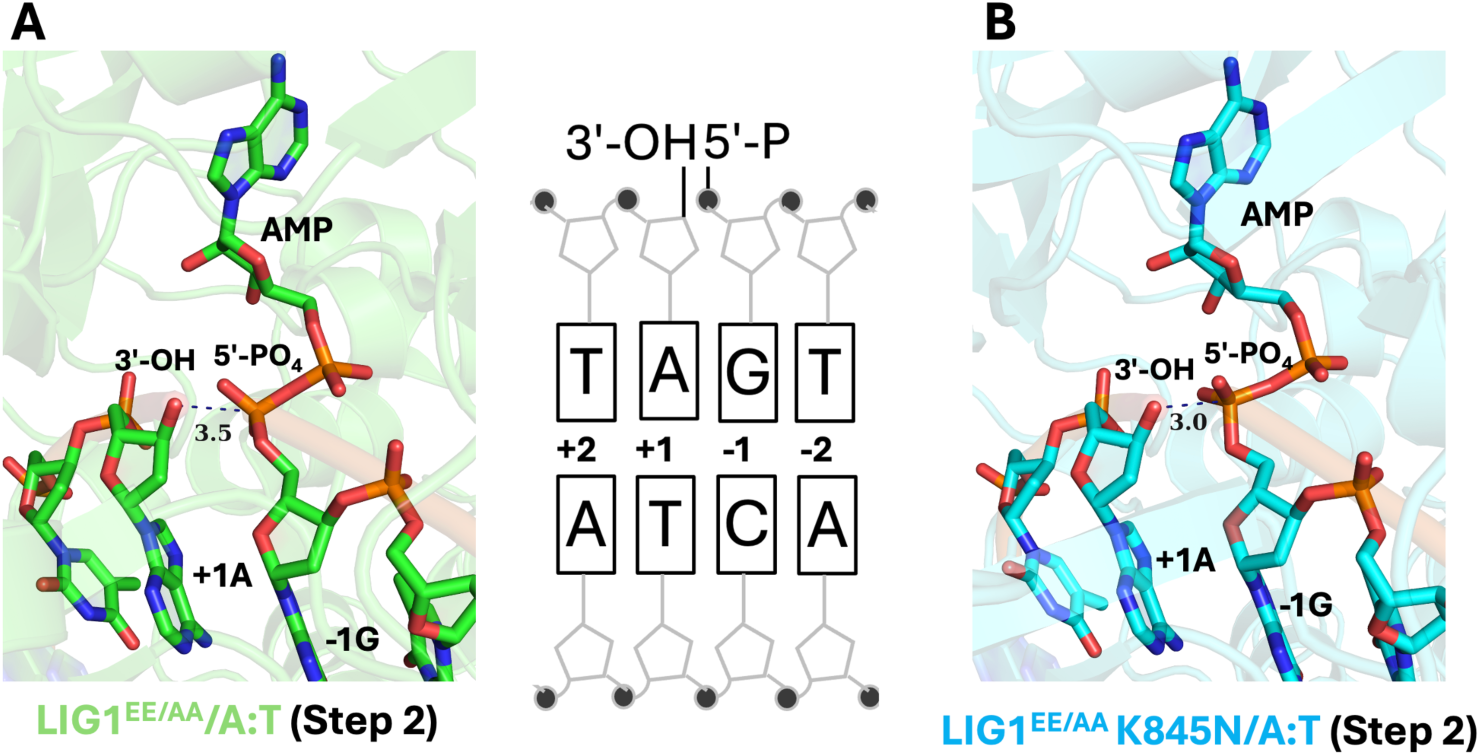
Structure of LIG1 K845N bound to the nick DNA duplex containing A:T at 3’-strand. **(A-B)** Structures of LIG1^EE/AA^ and LIG1^EE/AA^K845N in complex with nick DNA containing a cognate A:T show step 2 of the ligation reaction where AMP is bound to 5’-PO_4_ end of nick. The 2Fo -Fc density map of the AMP is contoured at 1.5σ (blue). LIG1 is shown in cartoon mode and DNA, AMP, active site residues are shown in stick mode. Scheme shows the sequence of nick DNA substrate used in LIG1 crystallization. LIG1^EE/AA^/3’-dA:T structure is previously solved (PDB: 7SUM).

**Figure 8.**
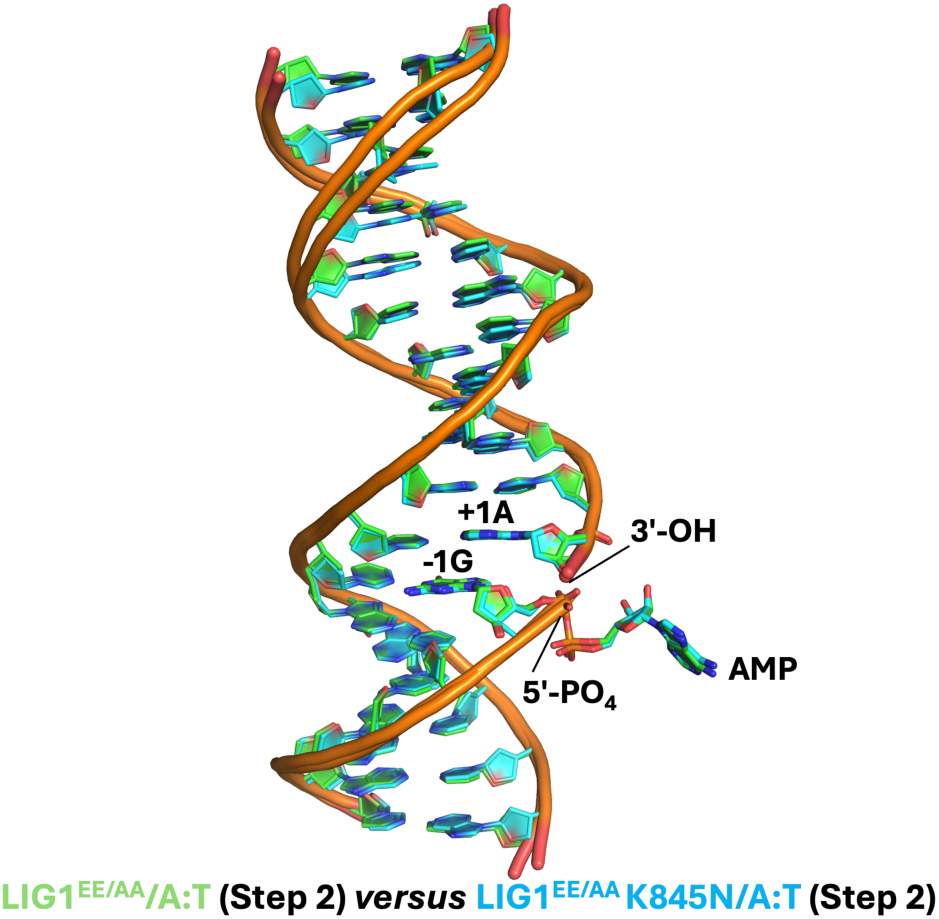
Superimposition of LIG1 structures. The overlay of LIG1^EE/AA^ and LIG1^EE/AA^K845N structures in complex with nick DNA containing canonical A:T shows similarity in the DNA conformation at the downstream of the nick.

We then further analyzed the impact of K845N mutation on the distances relative to the 3’-and 5’-ends of nick DNA (Figure 9). The distances between the functional groups of K845 and N845 side chains are calculated as 17.5Å and 15.8Å, respectively, relative to the 5’-end of nick. We observed 1Å difference in the distance between the functional groups of K845/N845 side chains with regards to the 3’-end of nick. These findings suggest that LIG1^EE/AA^ and LIG1^EE/AA^ K845N structures share similar configuration with the nick site containing canonical A:T end, and therefore, HD-associated amino acid substitution at K845 residue could not impact nick processing at the ligase active site. As we crystalized the ligase (LIG1^EE/AA^) in the absence of Mg^2+^, the metal ion cofactor and the low-fidelity background could impact structural arrangements at nick site when the ligation reaction proceeds. From the superimposition of the LIG1 structures (Supplementary Figure 11), we conclude that HD-associated variant K845N does not directly interact with the minor groove or template strand of the DNA as shown for previously solved LIG1 syndrome variants R641L and R771W (39). Furthermore, the ligation efficiency of LIG1 K845N *versus* these LIG1 syndrome variants demonstrated similar end joining ability of LIG1 K845N and P529L mutants, while nick sealing efficieny is compromised by LIG1 R641L and R771W mutations (Supplementary Figure 12).

**Figure 9.**
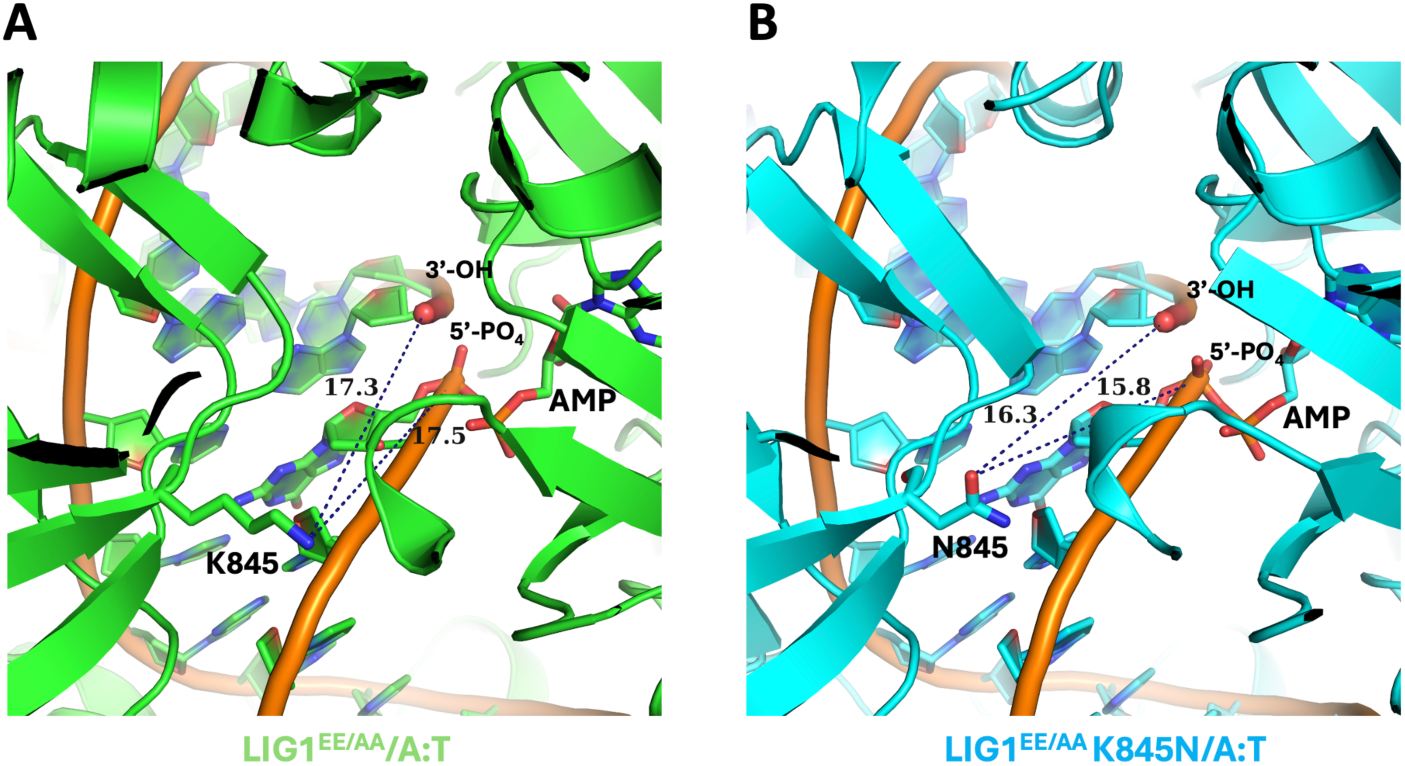
LIG1 structures demonstrate similarity in the distances relative to DNA ends. **(A-B)** Structures of LIG1^EE/AA^ and LIG1^EE/AA^K845N show the distances (>15Å) between the ligase active site residues K845 *versus* N845 relative to 3’-OH and 5’-PO_4_ ends of nick.

### Nick DNA binding by LIG1 K845N mutant at single-molecule level

Using TIRF microscopy, we employed single-molecule fluorescence co-localization approach to further investigate the impact of LIG1 K845N mutation on the nick DNA binding in real-time. For this purpose, we used the AF488-labeled dsDNA containing a single nick site with A:T end and Cy5-labeled LIG1 wild-type and K845N mutant proteins (Figure 10A).

**Figure 10.**
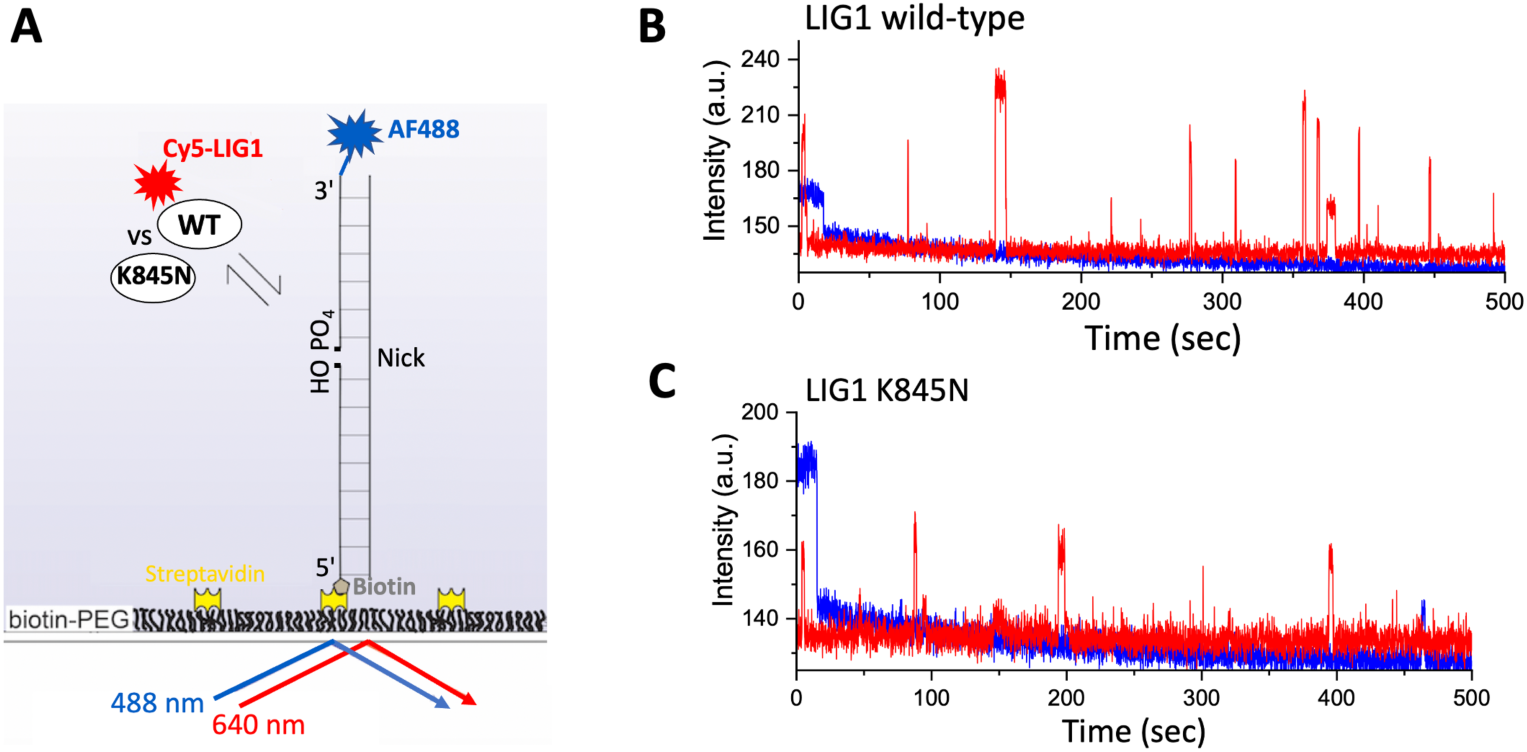
Single-molecule characterization of nick DNA binding by LIG1 K845N mutant. **(A)** Scheme shows that AF488-labeled dsDNA with a single nick site immobilized on PEG-coated, biotinylated slide surface for imaging with a TIRF microscope to monitor real-time Cy5-labeled LIG1/nick DNA binding. **(B-C)** Representative fluorescence intensity *versus* time traces show repeated protein binding events for LIG1 wild-type and K845N mutant.

LIG1^Cy5^ binding to nick DNA was identified by the co-localization of AF488 and Cy5 fluorescence signals within a diffraction-limited spot. The analyses of individual fluorescence time trajectories showed repeated transient Cy5 co-localizations with AF488 for wild-type and less frequent transient Cy5 co-localizations with AF488 for K845N mutant, indicating dynamic binding of LIG1 on the DNA (Figure 10B, C). Rastergrams generated from several individual time traces idealized by hidden Markov model (HMM) showed dynamic binding behavior for both LIG1 proteins to the nick DNA (Supplementary Figure 13).

From the dwell times distributions in the bound states, we next estimated the average bound and unbound lifetimes (Figure 11). For LIG1 wild-type, consistent with our recent study (22), we observed two populations with average binding lifetimes of 7.0 ± 0.9 sec (0.4 ± 0.1%) and 0.7 ± 0.1 sec (0.6 ± 0.1%). For LIG1 K845N mutant, the average binding lifetimes were calculated as 5.0 ± 0.2 sec (0.4 ± 0.01%) and 0.4 ± 0.1 sec (0.6 ± 0.01) as shown in differences in t_bound_ and t_unbound_ times (Figure 11A). We previously demonstrated that the long binding events for LIG1 and nick DNA interaction are due to the formation of stable nick-bound LIG1-DNA complex (22). When compared to LIG1 wild-type, K845N mutant binds less frequently as indicated by the increased t_unbound_ from 61 ± 2 sec (wild-type) to 100 ± 12 sec (K845N). The less frequent binding events as well as shorter binding lifetime time suggest that compared to LIG1 wild-type, the mutant has less affinity to find a target DNA, and also the stability of the nick DNA-LIG1 complex is relatively less (Figure 11B), suggesting that HD-associated mutation leads to a reduced affinity to bind to nick DNA.

**Figure 11.**
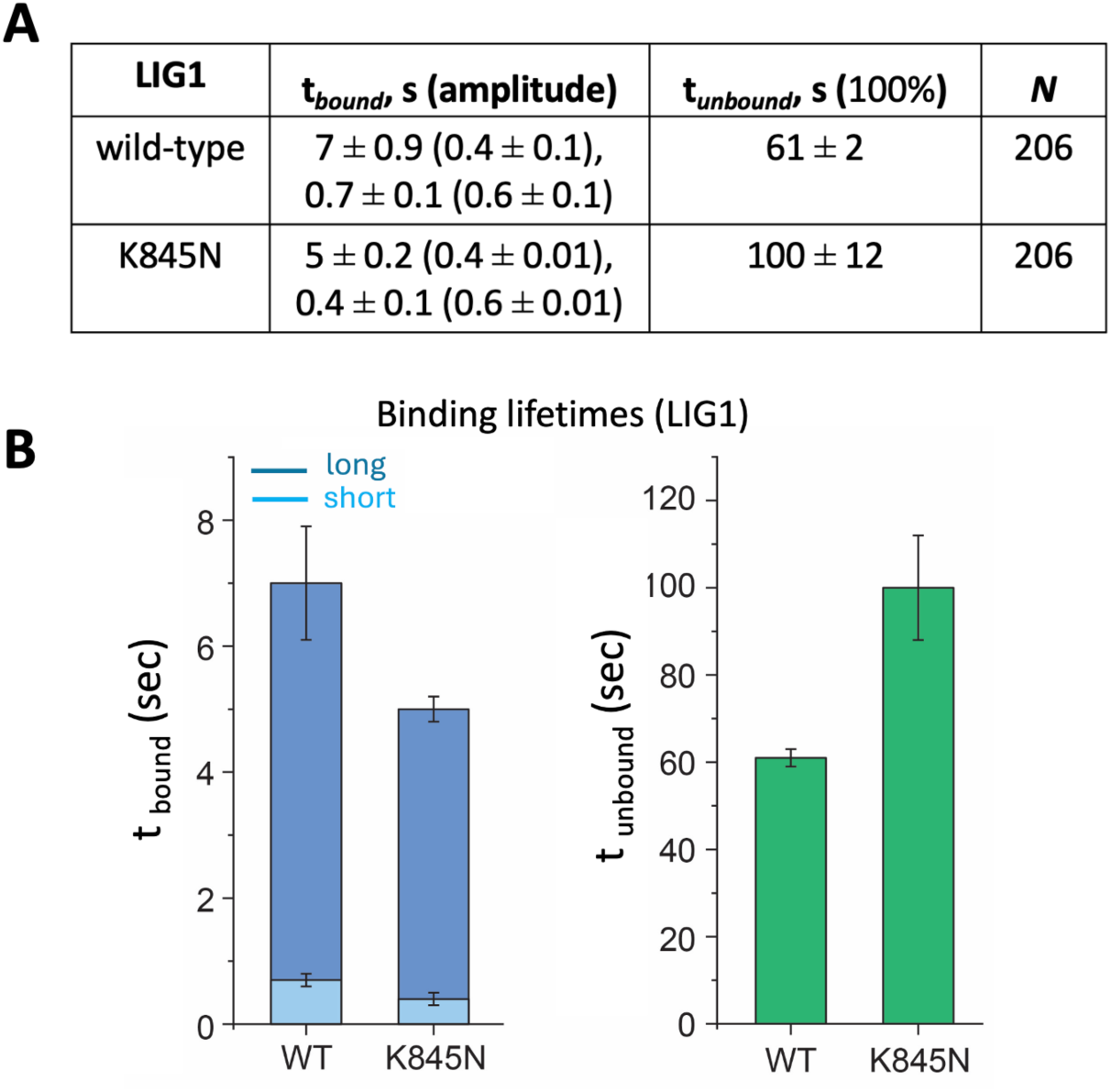
LIG1 K845N mutant binds less frequently with shorter binding lifetime to nick than wild-type. **(A-B)** Table shows the difference in lifetimes of protein-bound and -unbound states (A) and the bar graphs represents the comparison of *t*_bound_ and *t_un_*_bound_ times between LIG1 wild-type and K845N mutant (B).

## Discussion

Unstable CAG expansion in the coding region of the *HTT* gene leads to mHTT production, with eventual aggregation resulting in HD that causes nerve cells in the brain to decay over time (33–36). In the recent study (37), two alleles, CAA-loss and CAACAG-duplication mutants, were used to confirm the relationship between uninterrupted CAG repeat length and the disease onset, summarizing that the rate-driver of uninterrupted CAG repeats is independent of glutamine coding properties. This study also revealed that the mutation at K845 residue residing in the OBD domain of LIG1 is predicted to be onset delaying and suppresses CAG repeat expansion, the main determiner for the disease onset. LIG1, as the main replicative ligase, joins Okazaki fragments and seals final nick product at an ultimate step of DNA excision repair pathways (6). Yet, it remains unknown how the HD-associated hereditary LIG1 impairment due to K845N mutation could impact the function of the ligase at the final step. In the present study, we aim to comprehensively characterize the impact of LIG1 K845N variant on the ligase fidelity, DNA binding affinity, and nick sealing efficiency at biochemical, structural, and single-molecule levels.

Human immunodeficiency syndrome caused by the mutations in *LIG1* gene (P529L, R641L, R771W) leads to an inherited LIG1 deficiency, which was established from the individuals that exhibit growth retardation, developmental delays, sun sensitivity, and severe immunodeficiency, exhibits aberrant joining of Okazaki fragments and hypersensitivity to DNA alkylating agents (27–32). The amino acid residues that have been mutated in the LIG1 syndrome are located in the C-terminal catalytic domain of the protein, particularly in the AdD (E566), the DBD (P529) and the OBD (R771) domains (Supplementary Figure 14A). Our previous study demonstrated how these LIG1 variants affect the ligation of the DNA polymerase-promoted mutagenesis products (16). We reported that the LIG1 R641L and R771W mutants show subtle differences in the ligation efficiency for all possible 12 mismatches and a compromised mutagenic ligation of nick DNA with 8oxoG:A, while LIG1 P529L variant shows wild-type level of end joining ability to seal nicks with non-canonical ends.

Furthermore, X-ray structures of LIG1 variants R641L and R771W revealed a compromised cooperative network of interactions with the ligase active site, high-fidelity Mg^2+^ binding site, and DNA leading to a decrease in DNA binding affinity and ligation efficiency (39). Particularly, R641L leads to a disengagement in the DNA binding loop with connecting salt bridge to the active site (R641-D600 salt bridge anchor) as well as distortions in the DNA backbone, while LIG1 R771W causes remodeling of interdomain contacts at the juncture of the OBD and DBD domains, which is characterized by two salt bridges that support the OBD-DNA binding loop for stability in the minor groove. In their respective locations, both variants seem to disrupt the binding of protein-DNA-metal ion interactions that are usually involved in the mechanism by which LIG1 discriminates against nicks containing mismatches and damaged bases through Mg^2+^ binding mediated high-fidelity. From the overlay of previously solved structures of LIG1 syndrome variants R641L and R771W (39) *versus* HD-associated variant K845N we solved the structure for, the RMSD analysis suggest no significant changes in the overall structure (Supplementary Table 1). Furthermore, the superimposition of these structures also demonstrated that R641L and R771W variants interact with the minor groove of the DNA while K845N has no direct interaction with the DNA (Supplementary Figure 11). Our results from the ligation assays with a range of nick DNA substrates showed similar end joining ability of LIG1 wild-type in the presence of canonical ends, while we obtained a reduced ligation efficiency by LIG1 K845N mutant for all 12 possible non-canonical mismatches and 8oxoG. Regarding DNA-RNA substrates containing a single ribonucleotide at the 3’-end of nick, the ligation assays demonstrated a lack of sugar discrimination, suggesting that K845 side chain has no role for identification of “wrong” sugar at nick. Furthermore, the overlay of LIG1 structures in the presence and absence of N845 mutation demonstrated similar conformation and slight differences in the distances to both DNA ends. Further structural studies of LIG1 K845N mutant in complex with nicks containing mismatched or damaged ends could provide more insights into the mechanism by which the ligase active site carrying HD-associated K845N mutation ensures ligase fidelity. In addition, our single-molecule measurements in real-time revealed that LIG1 K845N variant binds to nick DNA less frequently for shorter life-times when compared to the wild-type enzyme, although both proteins can form a stable nick-bound LIG1-DNA complex. Because the conversion of nicks into phosphodiester bond requisites a rapid nick recognition to avoid deterioration of the exposed DNA ends to exonuclease degradation and increased frequency of recombination , this slower nick binding affinity we observed with LIG1 K845N variant may result in a delay in joining SSBs and the maturation of Okazaki fragments. In the event of DNA replication, this situation may cause replication forks to stall, activating fork rescue pathways that could involve recombination-based mechanisms that do not favor repeat expansion (40). It’s important to note a functional redundancy between LIG1 and LIG3α in DNA repair and replication (41). Furthermore, the recent studies reported back-up role of LIG3α in case of perturbation of the main DNA replication ligase, LIG1, leading to the formation of unligated Okazaki fragments during nuclear replication, which is trapped by Poly(ADP-Ribose) polymerase (42). Therefore, LIG3α may compensate this inefficiency caused by K845N mutation during DNA replication or repair leading to less reliance on LIG1.

The molecular basis underlying trinucleotide repeat expansions such as age-dependent neuronal CAG repeats has been found to be associated with the formation of non-B-form or DNA noncanonical structures that can compromise proper function of repair enzymes (43). This may cause to the integration of hairpin intermediates or loops with different size and stability into the genome (44). Further biochemical chatacterization of LIG1 K845N variant for processing of CAG repeat-containing substrates and coordination with other repair and replication proteins are required to explore mechanisms of repeat expansion as reported before for long-patch BER sub-pathway involving polβ-mediated strand displacement synthesis and FEN1 cleavage activities in case of strand slippage during processing of oxidative damage (45). Overall, our study contributes to understad how a defect in LIG1 ability to join strand breaks, stemming from a single K845N mutation, could lead to interruption in coordinated repair at the downstream steps during processing of a final nick product in normal *versus* disease states, underscoring an aberrant LIG1 function could play role in the pathogenesis of neurodegenerative diseases.

## Experimental Procedures

### Protein purifications

We generated Lys(K) to Asn(N) amino acid substitution at the 845 residue of LIG1 that resides in the Oligonucleotide binding domain (OBD) of the catalytic core (Supplementary Figure 14A). We generated the K845N mutation in the LIG1 C-terminal (△261) and full-length (1-919 amino acids) background (pET-24b) of LIG1. The recombinant proteins wild-type and K845N mutant with 6x his-tag for LIG1 C-terminal and full-length proteins were purified as described previously (12–22). Briefly, the proteins were overexpressed in *E. coli* (DE3) cells and grown in Terrific Broth (TB) media with kanamycin (50 μgml^−1^) and chloramphenicol (34 μgml^−1^) at 37 °C. Once the OD_600_ reached 1.0, cells were induced with 0.5 mM isopropyl β-D-thiogalactoside (IPTG) and overexpression continued overnight at 20 °C. After centrifugation, cells were lysed in the lysis buffer containing 50 mM Tris-HCl (pH 7.0), 500 mM NaCl, 20 mM imidazole, 2 mM β-mercaptoethanol, 5% glycerol, and 1 mM phenylmethylsulfonyl fluoride (PMSF) by sonication at 4 °C. The cell lysate was pelleted at 31, 000 x g for 90 min at 4 °C. The cell lysis solution was clarified and then loaded onto a HisTrap HP column that was previously equilibrated with the binding buffer containing 50 mM Tris-HCl (pH 7.0), 500 mM NaCl, 20 mM imidazole, 2 mM β-mercaptoethanol, and 5% glycerol. The column was washed with the binding buffer and then eluted with an increasing imidazole gradient 20-500 mM at 4 °C. The collected fractions were then subsequently loaded onto a HiTrap Heparin column that was equilibrated with the binding buffer containing 20 mM Tris-HCl (pH 7.0), 50 mM NaCl, 2 mM β-mercaptoethanol, and 5% glycerol. The protein was eluted with a linear gradient of NaCl up to 1 M. LIG1 proteins were further purified by a Superdex 200 10/300 column in the buffer containing 50 mM Tris-HCl (pH 7.0), 200 mM NaCl, and 1 mM DTT, and 5% glycerol. The LIG1 proteins purified in this study were concentrated, aliquoted, and stored at -80 °C. Protein quality was evaluated on a 10% SDS-PAGE gel, and protein concentrations were measured using absorbance at 280 nm (Supplementary Figure 14B).

### DNA ligation assays

Nick DNA substrates with a 6-carboxyfluorescein (FAM) label were used in DNA ligation assays. Nick DNA substrates containing Watson-Crick base paired ends, and all 12 non-canonical mismatches were prepared by annealing upstream oligonucleotides 3’-dA, dT, dG, or dC with template oligonucleotides A, T, G, or C (Supplementary Table 2). Nick DNA substrates containing damaged ends were prepared by annealing 3’-8oxodG or 3’-8oxorG with template oligonucleotides A or C (Supplementary Table 3). Nick DNA substrates containing a single ribonucleotide at the 3’-end were prepared by annealing upstream oligonucleotides 3’-rA, 3’-rG, and 3’-rC with template oligonucleotides A, T, G, or C (Supplementary Table 4). DNA ligation assays were performed as described (12–22) to investigate the substrate specificity of LIG1 wild-type and K845N mutant for nick DNA substrates containing canonical, mismatched, damaged, and ribonucleotide-containing ends (Supplementary Scheme 1). The reaction was initiated by the addition of LIG1 (100 nM) to a mixture containing 50 mM Tris-HCl (pH 7.5), 100 mM KCl, 10 mM MgCl_2_, 1 mM ATP, 1 mM DTT, 100 µgml^-1^ BSA, 1% glycerol, and the nick DNA substrate (500 nM) in a final volume of 10 µl. The reaction mixture was incubated at 37 °C and stopped at the time points indicated in the figure legends by mixing with an equal volume of loading dye containing 95% formamide, 20 mM ethylenediaminetetraacetic acid, 0.02% bromophenol blue, and 0.02% xylene cyanol. Reaction products were separated by electrophoresis on an 18% Urea-PAGE gel, the gels were scanned with the Typhoon PhosphorImager RGB, and the data was analyzed using ImageQuant software.

### Crystallization and structure determination

For the crystallization of LIG1 K845N, we used LIG1 C-terminal (△261) and the EE/AA mutant that harbors E346A and E592A mutations, resulting in the ablation of the high-fidelity site, which has been utilized in our previous LIG1 structures with non-canonical nicks (23–26). We solved the structure of LIG1 K845N in complex with nick DNA containing a canonical (A:T) end (Supplementary Table 5). LIG1 (at 27 mgml^-1^)/DNA complex solution was prepared in the buffer containing 20 mM Tris-HCl (pH 7.0), 200 mM NaCl, 1 mM DTT, 1 mM EDTA, and 1 mM ATP at 1.4:1 DNA:protein molar ratio and then mixed with 1 μl reservoir solution. LIG1-nick DNA complex crystals were grown at 20 °C using the hanging drop method, harvested, and submerged in cryoprotectant solution containing reservoir solution mixed with glycerol to a final concentration of 20% glycerol before being flash cooled in liquid nitrogen. LIG1 crystals were obtained and kept at 100 °K during X-ray diffraction data collection using the beamline CHESS-7B2. The collected data were reduced and scaled using HKL2000 (HKL Research, Inc). LIG1 K845N structure was solved by the molecular replacement method using PHASER with PDB entry 7SUM as a search model (46). Iterative rounds of model building were performed in COOT and the final models were refined with PHENIX (47, 48). All structural images were drawn using PyMOL (The PyMOL Molecular Graphics System, V0.99, Schrödinger, LLC). Detailed crystallographic statistics are provided in Table 1.

### Single-molecule nick DNA binding measurements

We performed single-molecule imaging experiments to monitor LIG1-nick DNA binding using a total internal reflection fluorescence (TIRF) microscope (Nikon Eclipse Ti2-E) as reported previously (22). For this purpose, we fluorescently labeled LIG1 full-length proteins (wild-type and K845N mutant) by Amersham Cy5 monoreactive NHS ester dye (LIG1^Cy5^) and pre-adenylated the LIG1^Cy5^ proteins. We used 3’-AF488-labeled dsDNA substrate (34-mer) including 5’-Biotin and a single nick site containing an A:T end (Supplementary Table 6). Glass coverslips were functionalized with a mixture of biotin-PEG-SVA and mPEG-SVA and the microfluidic channels were assembled using the passivated coverslips and Grace Bio-Labs HybriWell™ sealing system, which were rinsed three times with the T50 buffer containing 10 mM Tris-HCl (pH 8.0) and 50 mM NaCl. Streptavidin (0.2 mg/ml) in the T50 buffer was then flowed onto the slide, reacted with the biotin-PEG, and then washed with the T50 buffer. The DNA substrate was diluted to a final concentration of 10 pM in the imaging buffer containing 1 mM HEPES (pH 7.4), 20 mM NaCl, 0.02% BSA (w/v), and 0.002% Tween 20 (v/v), flowed onto the slide, and an excess unbound DNA substrate was then washed by flowing with the imaging buffer. Oxygen-scavenging system (OSS) consisting of 44 mM glucose, 165 U/ml glucose oxidase from Aspergillus niger and catalase (2, 170 U/ml) were added to slow photobleaching. 10 mM Trolox was added to reduce photo blinking. Finally, LIG1^Cy5^ proteins (1 nM) in the imaging buffer containing the OSS mixture were flowed onto the slide and allowed to equilibrate before imaging with an objective-based TIRF microscope. Both AF488 and Cy5 dyes were simultaneously excited using 488 nm and 640 nm lasers (power 5 mW), respectively. Emissions from two fluorophores were separated into two channels using a Cairn Optosplit II image splitter and simultaneously recorded at 100 ms time resolution using a Hamamatsu SCMOS camera (77054115) using NIS-Elements acquisition software (Nikon, version: AR 6.02.01). The locations of molecules and fluorophore intensity over time traces were extracted from the raw movie files using Nikon NIS-Elements analysis software (Nikon, version: AR 6.02.01). Genuine fluorescence time traces for individual molecules were selected using NIS-Elements time measurement analysis and were idealized using a two-state hidden Markov model (HMM) for the unbound and bound states in QuB. Rastergrams summarizing several individual traces were generated from the individual trace HMMs using a custom written MATLAB script. From the idealized traces, dwell times of the bound and unbound states were calculated using MATLAB. Cumulative frequency of the bound and unbound dwell-time distributions was plotted and fitted in Origin Lab (version 2024b) with single or double exponential functions to obtain the bound (t_bound_) and unbound (t_unbound_) state lifetimes.

### Data Availability

Correspondence and requests for materials should be addressed to Melike Çağlayan. Atomic coordinates and structure factors for the reported crystal structures of LIG1 K845N have been deposited in the RCSB Protein Data Bank under accession number of 9NYS. All relevant data are available from the authors upon reasonable request.

## Supporting information

Manuscript

## Acknowledgement

This work is based upon research conducted at the Center for High Energy X-ray Sciences (CHEXS), which is supported by the National Science Foundation under award DMR-1829070, and the Macromolecular Diffraction at CHESS (MacCHESS) facility, which is supported by award 1-P30-GM124166-01A1 from the National Institute of General Medical Sciences, National Institutes of Health, and by New York State’s Empire State Development Corporation (NYSTAR).

## Funding

This work was supported by a grant 1R35GM147111-01 from the National Institute of General Medical Sciences (NIGMS) to M.Ç.

## Conflict of interest

The authors declare that they have no conflicts of interest with the contents of this article.

## Author Contributions

M.Ç.: conceptualization; J.R., K.L.: purified the proteins and performed biochemical experiments. K.B.: solved LIG1 structures. S.C.: performed single-molecule experiments. J.R., K.L., S.C.: writing-reviewing. M.Ç.: supervision, writing-reviewing, revisions, and editing. M.Ç.: funding.

## Notes

### Competing Interest Statement

The authors have declared no competing interest.

