## Supplementary material for "Characterization of nick binding and sealing by LIG1 Huntington’s disease-asssociated K845N variant at biochemical, structural, and single-molecule levels": Manuscript

### Supplementary Information

Supplementary Figures 1-14

Supplementary Scheme 1

Supplementary Tables 1-6

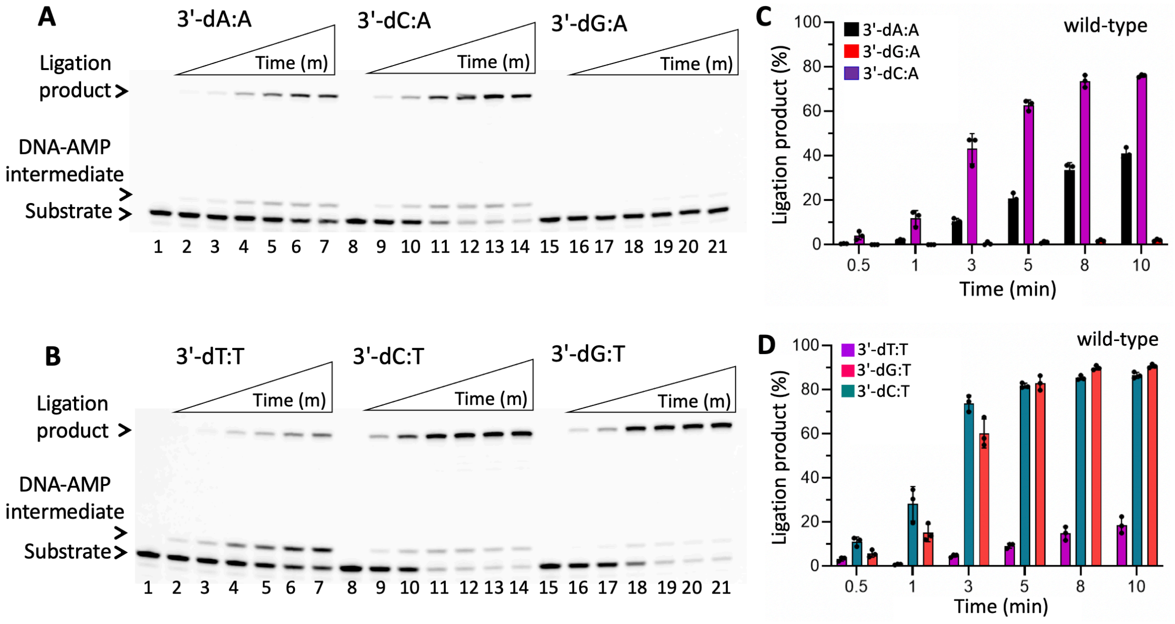


**Supplementary Figure 1. Ligation efficiency of LIG1 wild-type for nick DNA substrates containing template A and T mismatches. (A)** Lanes 1, 8, and 15 are the negative enzyme controls of the nick DNA substrates with 3'-dA:A, 3'-dC:A, and 3'-dG:A mismatches, respectively. **(B)** Lanes 1, 8, and 15 are the negative enzyme controls of the nick DNA substrates with 3'-dT:T, 3'-dC:T, and 3'-dG:T mismatches, respectively. In both panels, lanes 2-7, 9-14, and 16-21 are the ligation reaction products by LIG1 wild-type, and correspond to time points of 0.5, 1, 3, 5, 8, and 10 min. **(C-D)** Graphs show time-dependent changes in the amount of ligation products. The data represent the average from three independent experiments ± SD.

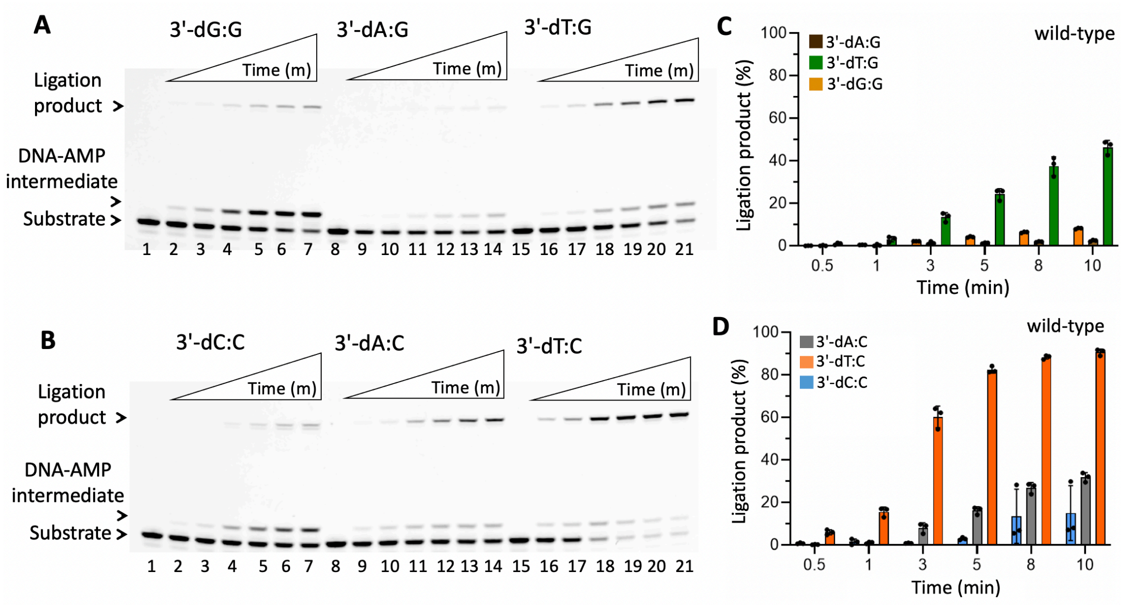


**Supplementary Figure 2. Ligation efficiency of LIG1 wild-type for nick DNA substrates containing template G and C mismatches. (A)** Lanes 1, 8, and 15 are the negative enzyme controls of the nick DNA substrates with 3'-dG:G, 3'-dA:G, and 3'-dT:G mismatches, respectively. **(B)** Lanes 1, 8, and 15 are the negative enzyme controls of the nick DNA substrates with 3'-dC:C, 3'-dA:C, and 3'-dT:C mismatches, respectively. In both panels, lanes 2-7, 9-14, and 16-21 are the ligation reaction products by LIG1 wild-type, and correspond to time points of 0.5, 1, 3, 5, 8, and 10 min. **(C-D)** Graphs show time-dependent changes in the amount of ligation products. The data represent the average from three independent experiments ± SD.

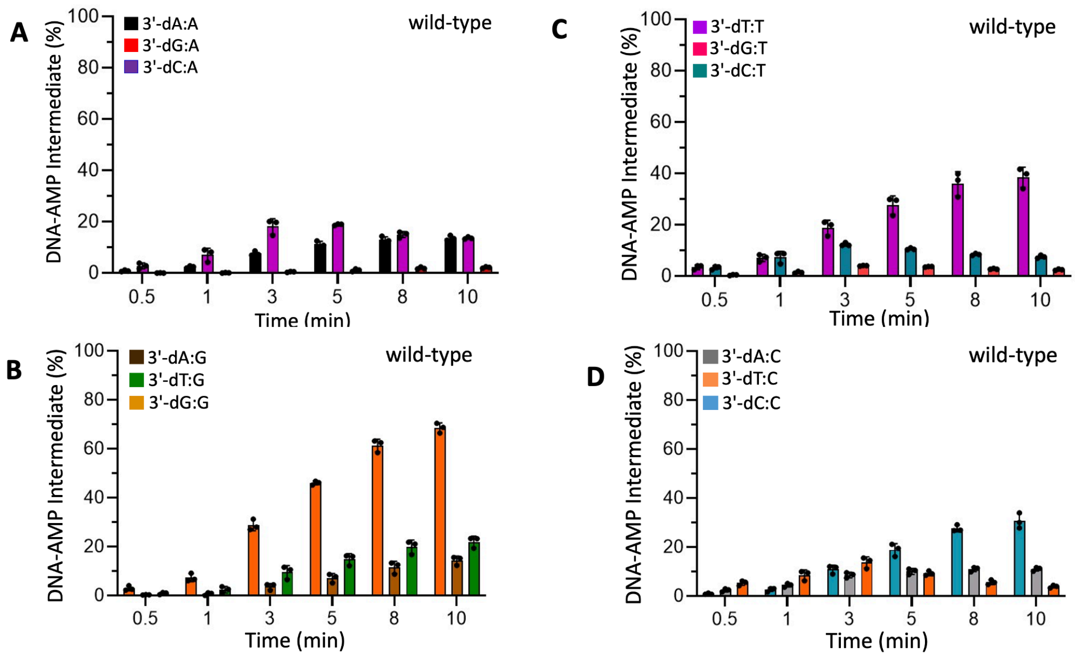


**Supplementary Figure 3. DNA-AMP intermediate formation during ligation of nick substrates containing template A, T, G, and C mismatches by LIG1 wild-type. (A-D)** Graphs show time-dependent changes in the amount of DNA-AMP intermediate products. The data represent the average from three independent experiments ± SD.

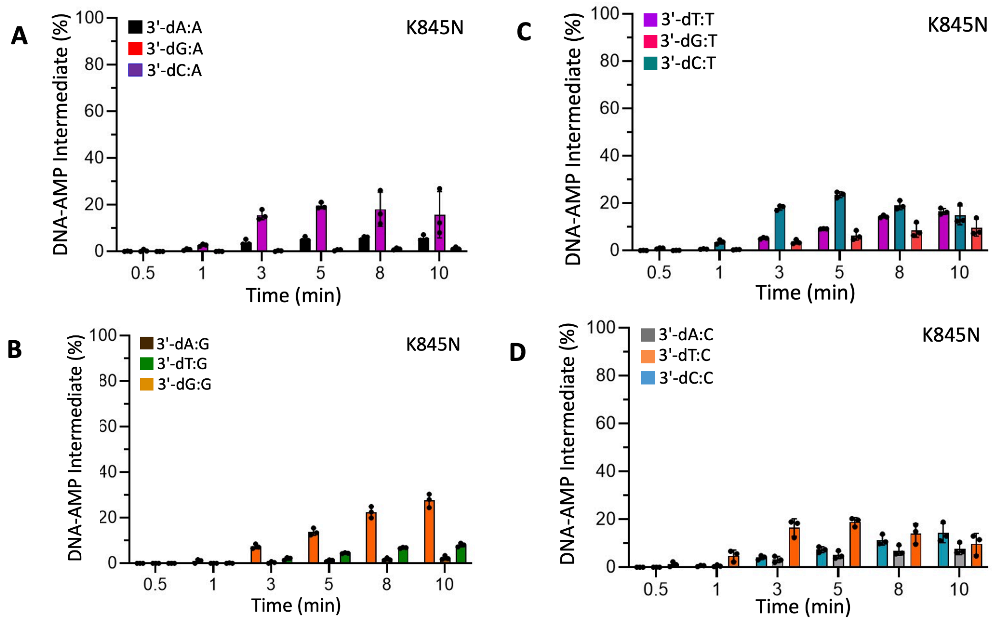


**Supplementary Figure 4. DNA-AMP intermediate formation during ligation of nick substrates containing template A, T, G, and C mismatches by LIG1 K845N mutant. (A-D)** Graphs show time-dependent changes in the amount of DNA-AMP intermediate products. The data represent the average from three independent experiments ± SD.

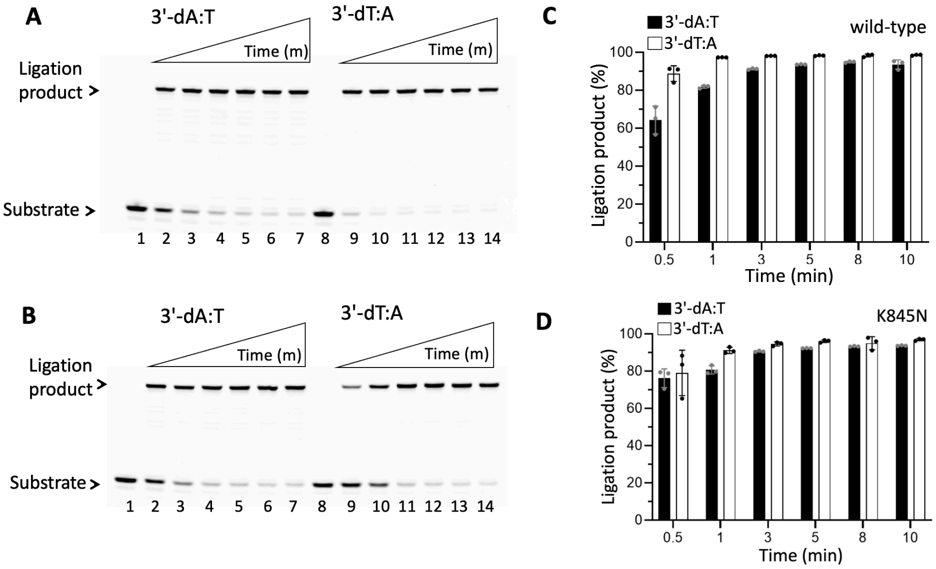


**Supplementary Figure 5. Ligation efficiency of LIG1 wild-type *versus* K845N mutant for nick DNA substrates containing canonical ends. (A-B)** Lanes 1 and 8 are the negative enzyme controls of the nick DNA substrates with 3'-dA:T and 3'-dT:A respectively. In both panels, lanes 2-7 and 9-14 are the ligation reaction products by LIG1 wild-type (A) and K845N mutant (B), and correspond to time points of 0.5, 1, 3, 5, 8, and 10 min. **(C-D)** Graphs show time-dependent changes in the amount of ligation products. The data represent the average from three independent experiments ± SD.

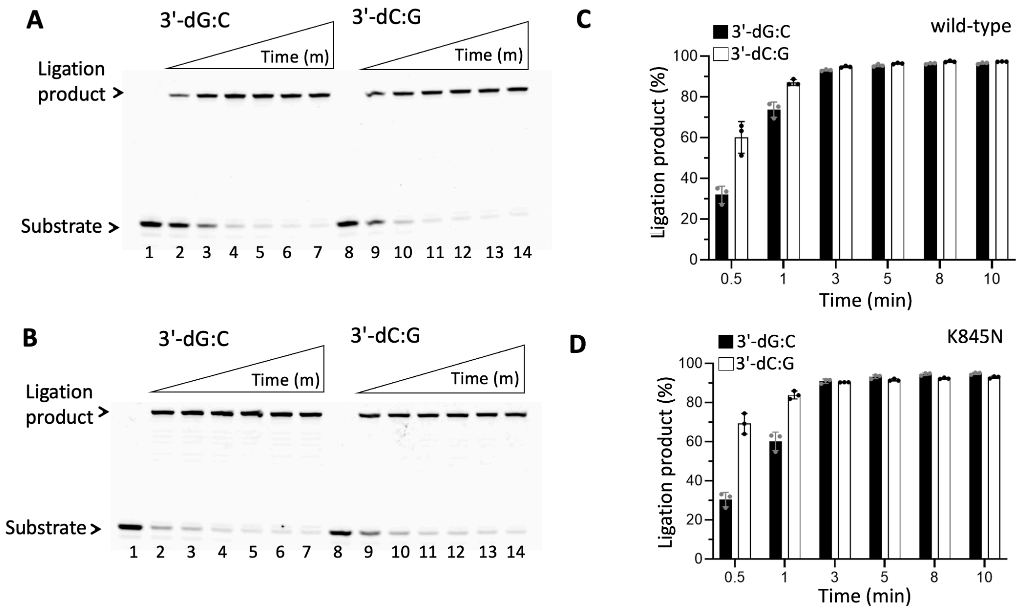


**Supplementary Figure 6. Ligation efficiency of LIG1 wild-type *versus* K845N mutant for nick DNA substrates containing canonical ends. (A-B)** Lanes 1 and 8 are the negative enzyme controls of the nick DNA substrates with 3'-dG:C and 3'-dC:G respectively. In both panels, lanes 2-7 and 9-14 are the ligation reaction products by LIG1 wild-type (A) and K845N mutant (B), and correspond to time points of 0.5, 1, 3, 5, 8, and 10 min. **(C-D)** Graphs show time-dependent changes in the amount of ligation products. The data represent the average from three independent experiments ± SD.

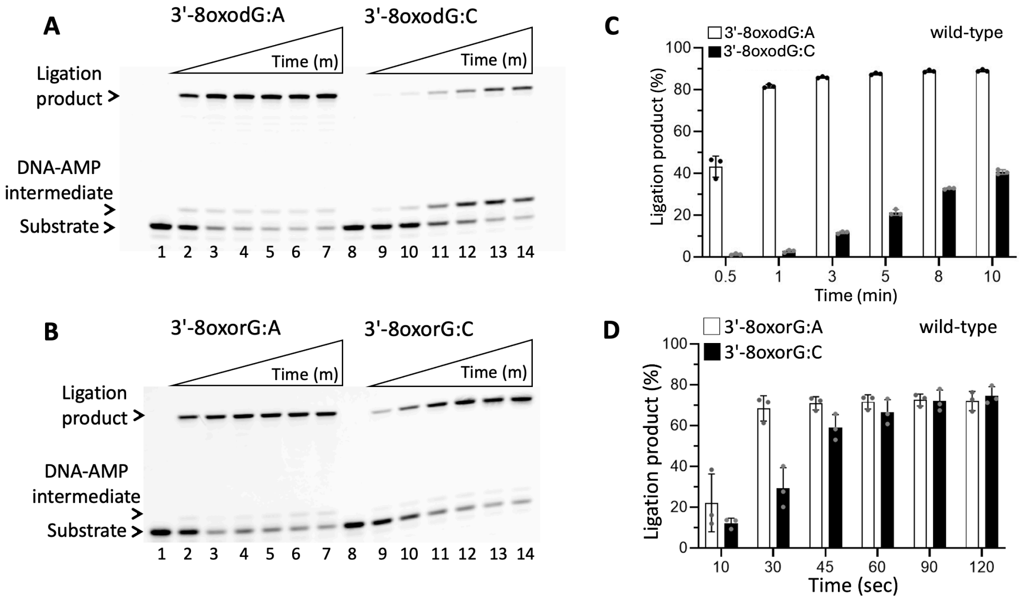


**Supplementary Figure 7. Ligation efficiency of LIG1 wild-type for nick DNA substrates containing oxidatively damaged ends. (A,C)** Lanes 1 amd 8 are the negative enzyme controls of the nick DNA substrates with 3'-8oxodG:A and 3'-8oxodG:C, respectively. Lanes 2-7 and 9-14 are the ligation reaction products by LIG1 wild-type, and correspond to time points of 0.5, 1, 3, 5, 8, and 10 min. Graph shows time-dependent changes in the amount of ligation products. The data represent the average from three independent experiments ± SD. **(B,D)** Lanes 1 and 8 are the negative enzyme controls of the nick DNA substrates with 3'-8oxorG:A and 3'-8oxorG:C, respectively. Lanes 2-7 and 9-14 are the ligation reaction products by LIG1 wild-type, and correspond to time points of 0.5, 1, 3, 5, 8, and 10 min. Graph shows time-dependent changes in the amount of ligation products. The data represent the average from three independent experiments ± SD.

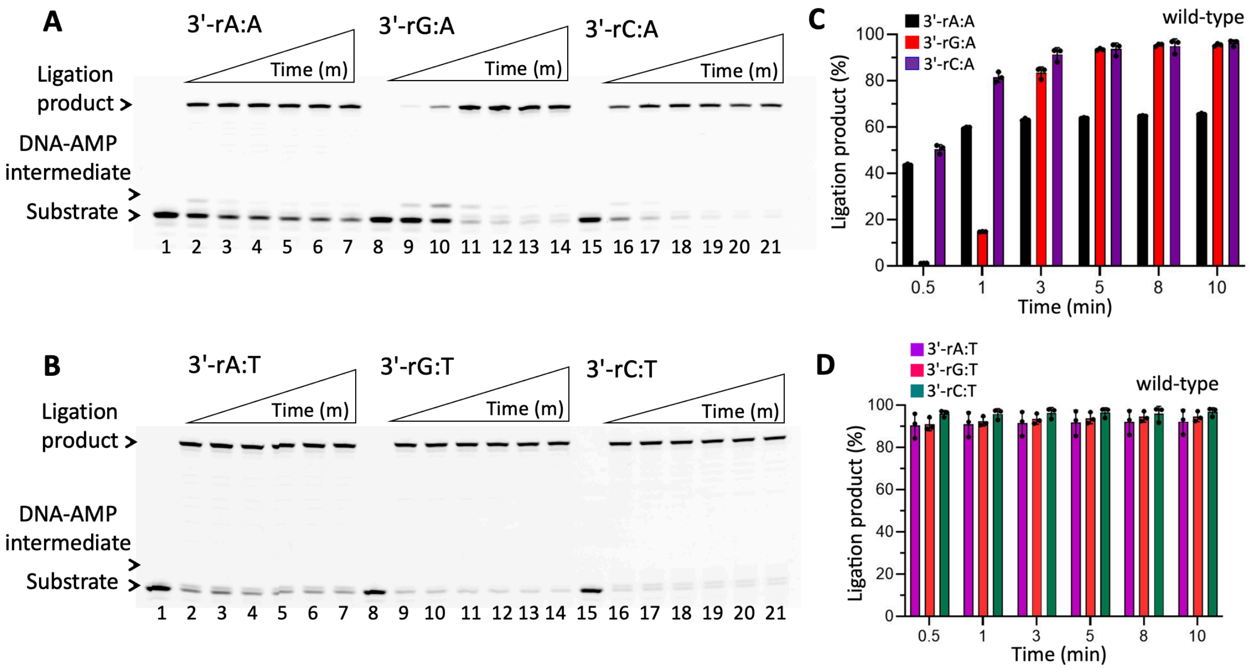


**Supplementary Figure 8. Ligation efficiency of LIG1 wild-type for nick DNA substrates containing 3'-ribonucleotides opposite to template A and T. (A)** Lanes 1, 8, and 15 are the negative enzyme controls of the nick DNA substrates with 3'-rA:A, 3'-rG:A, and 3'-rC:A, respectively. **(B)** Lanes 1, 8, and 15 are the negative enzyme controls of the nick DNA substrates with 3'-rA:T, 3'-rG:T, and 3'-rC:T, respectively. In both panels, lanes 2-7, 9-14, and 16-21 are the ligation reaction products by LIG1 wild-type, and correspond to time points of 0.5, 1, 3, 5, 8, and 10 min. **(C-D)** Graphs show time-dependent changes in the amount of ligation products. The data represent the average from three independent experiments ± SD.

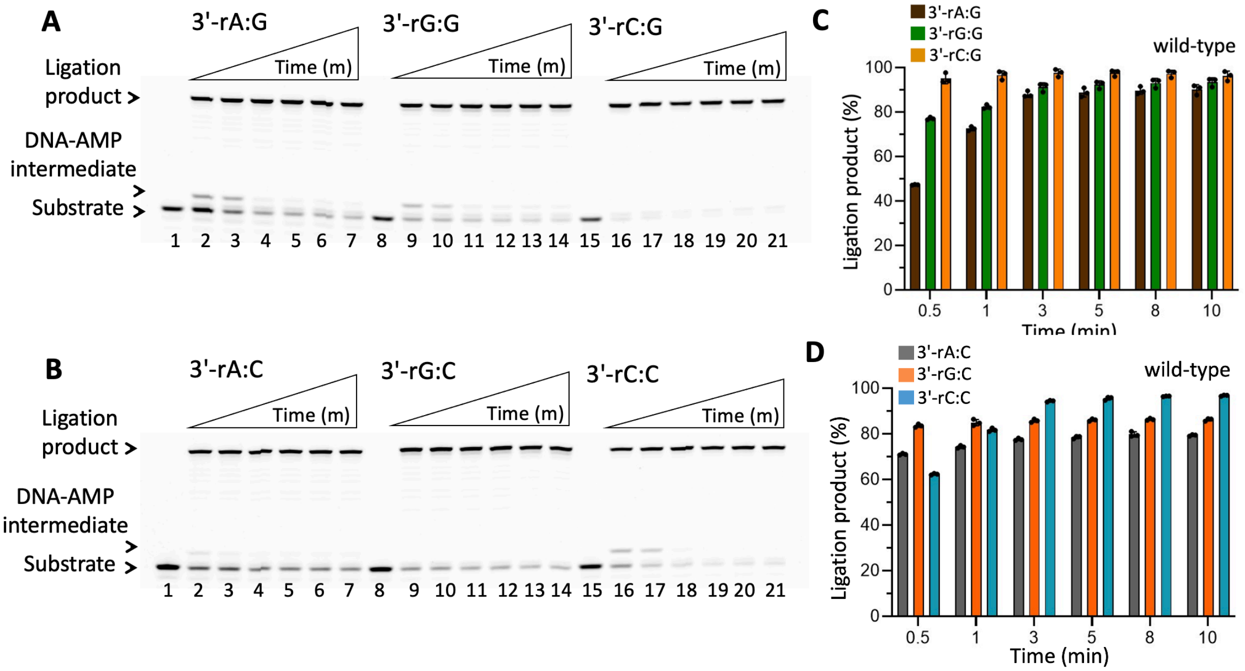


**Supplementary Figure 9. Ligation efficiency of LIG1 wild-type for nick DNA substrates containing 3'-ribonucleotides opposite to template G and C. (A)** Lanes 1, 8, and 15 are the negative enzyme controls of the nick DNA substrates with 3'-rA:G, 3'-rG:G, and 3'-rC:G, respectively. **(B)** Lanes 1, 8, and 15 are the negative enzyme controls of the nick DNA substrates with 3'-rA:C, 3'-rG:C, and 3'-rC:C, respectively. In both panels, lanes 2-7, 9-14, and 16-21 are the ligation reaction products by LIG1 wild-type, and correspond to time points of 0.5, 1, 3, 5, 8, and 10 min. **(C-D)** Graphs show time-dependent changes in the amount of ligation products. The data represent the average from three independent experiments ± SD.

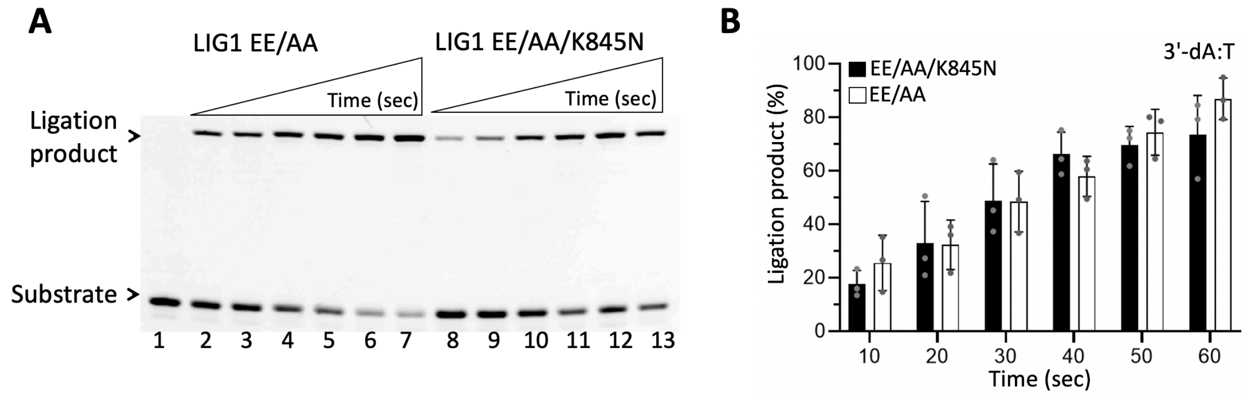


**Supplementary Figure 10. Ligation efficiency of LIG1 K845N mutant in the low-fidelity background. (A)** Line 1 is the negative enzyme control of the nick DNA substrate with 3'-dA:T. Lanes 2-7 and 8-13 are the ligation reaction products by LIG1 EE/AA low-fidelity double mutant and LIG1 EE/AA/K845N triple-mutant, respectively, and correspond to time points of 10, 20, 30, 40, 50, and 60 sec. **(B)** Graph shows time-dependent changes in the amount of ligation products. The data represent the average from three independent experiments ± SD.

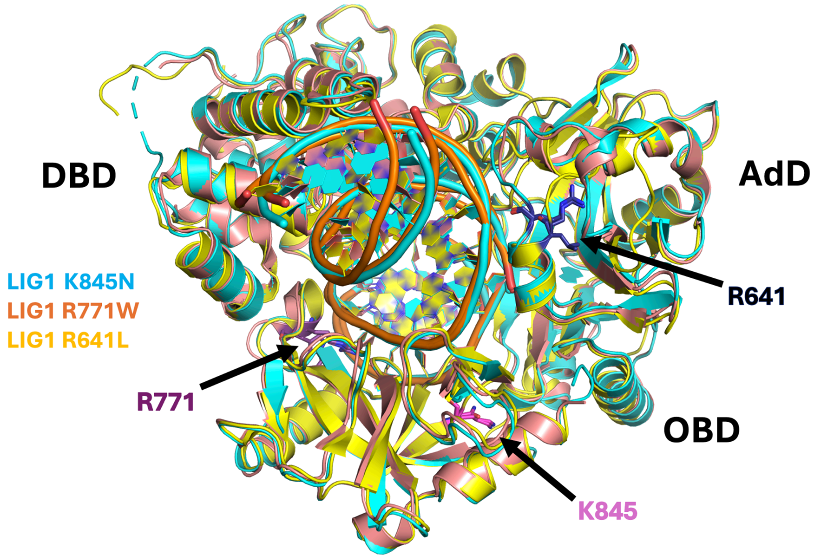


**Supplementary Figure 11. The superimposition of LIG1 disease mutant structures.** The overlay of LIG1 structures for LIG1 syndrome mutants R641L (blue) and R771W (purple) with HD-associated variant K845N (cyan) in the presence of nick containing canonical end reveals that the LIG1 syndrome mutants directly interacts with the minor groove/template strand of the DNA, while the K845N does not interact with the minor or major groove of the DNA. LIG1 syndrome mutations are located in the AdD (R641L) and the OBD (R771W) domains, while LIG1 HD-asssociated K845N mutant resides in OBD domain of LIG1. The structures of LIG1 syndrome mutants were previously solved by other group (x) for LIG1$^{R641L}$ (7L34) and LIG1$^{R771W}$ (7L35).

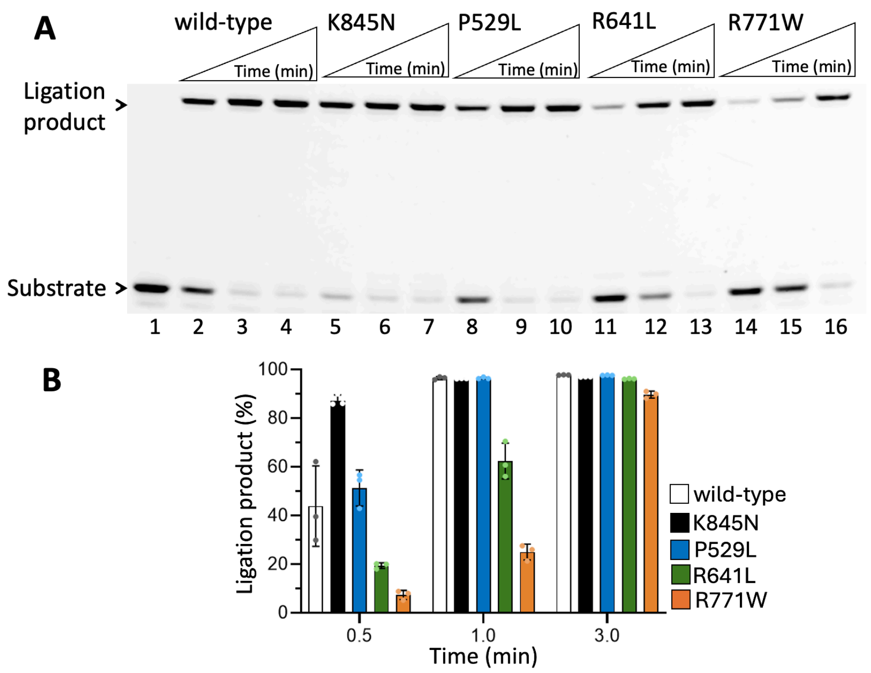


**Supplementary Figure 12. The comparision of ligation efficiency by all LIG1 disease variants. (A)** Line 1 is the negative enzyme control of nick substrate with 3'-dA:T. Lanes 2-4, 5-7, 8-10, 11-13, and 14-16 are the ligation reaction products by LIG1 EE/AA low-fidelity mutant, LIG1 wild-type, and LIG1 syndrome mutants P529L, R641L, R771W, and LIG1 HD-associated variant K845N, and correspond to time points of 0.5, 1, 3, 5, 8, and 10 min. **(B)** Graph shows time-dependent changes in the amount of ligation products. The data represent the average from three independent experiments ± SD.

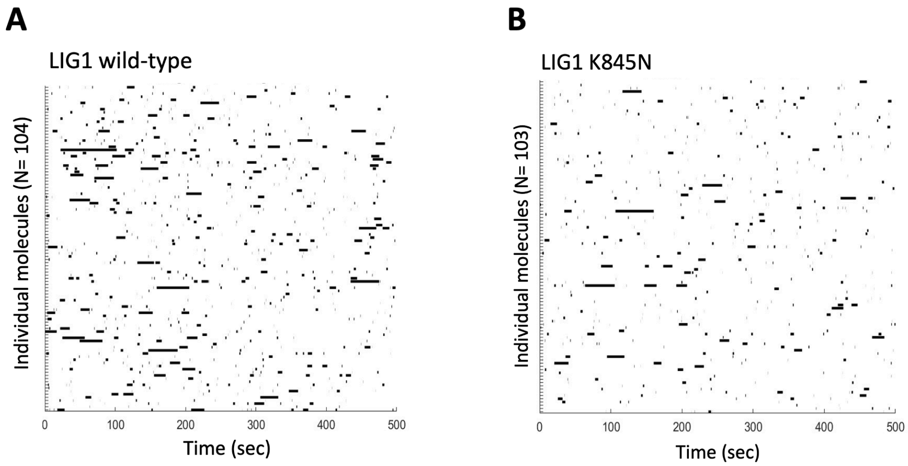


**Supplementary Figure 13. Single-molecule nick DNA binding of LIG1. (A-B)** Rastergrams of randomly selected traces are shown for LIG1 proteins displaying the distinct nick DNA binding behavior for wild-type and K845N mutant.

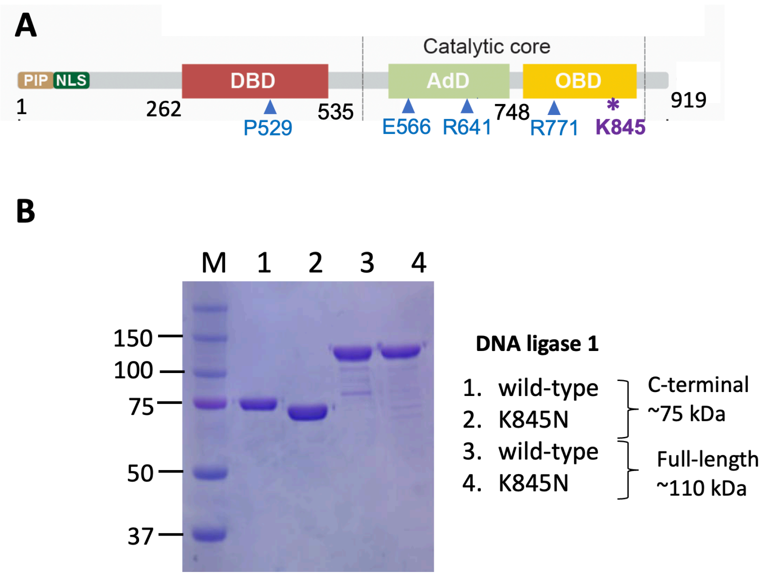


**Supplementary Figure 14. Purified LIG1 proteins used in the study. (A)** The domain organization of DNA ligase 1 (LIG1) protein (1-919 amino acids) including N-terminal region (1-262 amino acids) and C-terminal domain that contains DNA-binding domain (DBD) and the catalytic core consisting of Adenylation (AdD) and Oligonucleotide-binding (OBD) domains. The mutations at the amino acid residues that have been associated with LIG1 syndrome are located in the C-terminal catalytic domain of the protein, particularly in the DBD (P529), the AdD (E566, R641), and the OBD (R771) domains. LIG1 HD disease-asssociated mutation K845N reside in OBD domain of the catalytic core. **(B)** Final purity of LIG1 C-terminal and full length proteins used in this study. M represents the protein marker.

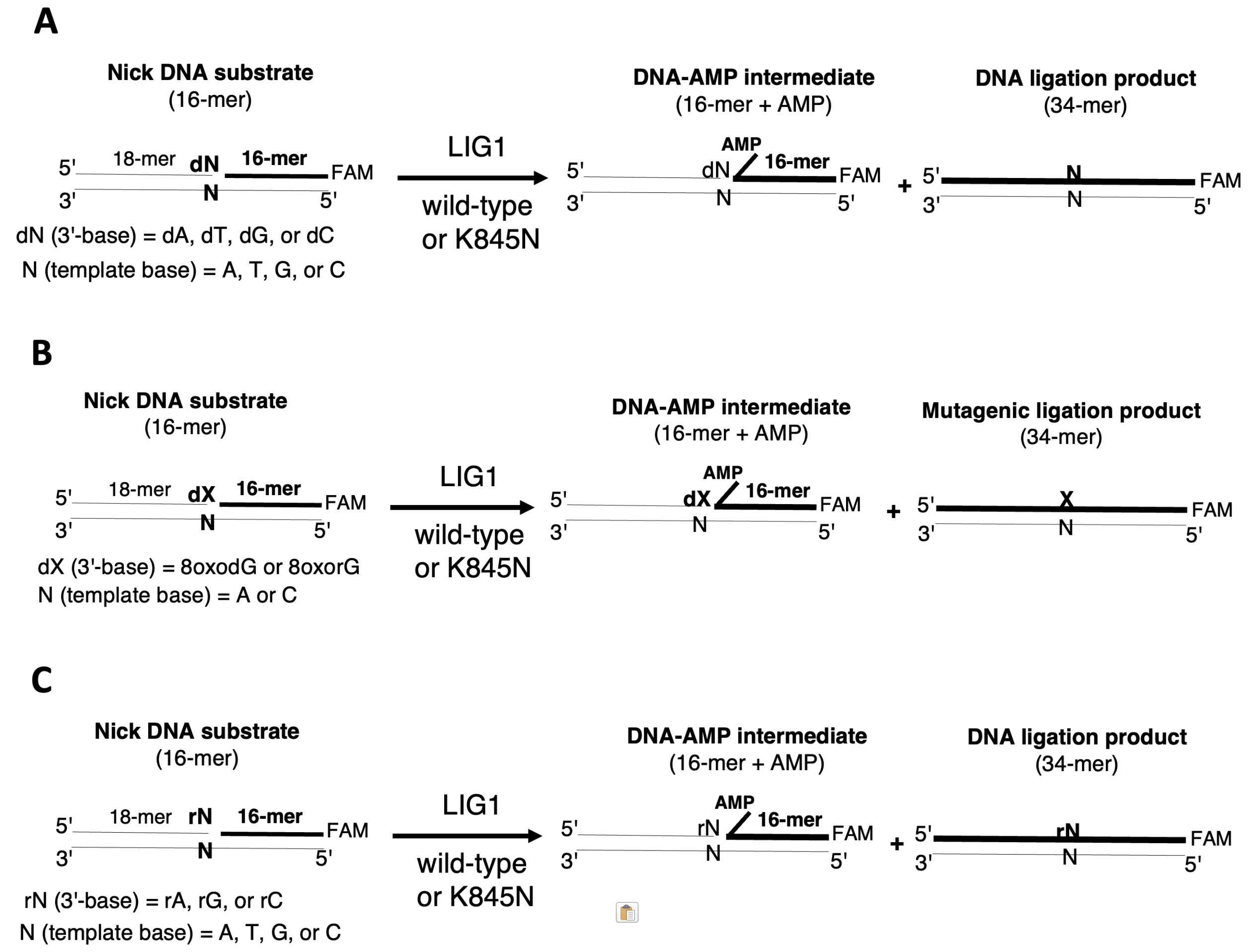


**Supplementary Scheme 1. Illustrations of DNA ligation assays used in this study. (A-C)** Ligation assays were used to evaluate the substrate specificity of LIG1 wild-type and K845N mutant for the nick DNA substrates including 3'-mismatches (A), 3'-8oxodG or 3'-8oxorG (B), and 3'-ribonucleotide (C). Reaction products include nick sealing and DNA-AMP intermediate with 5'-adenylate (AMP).

| RMSD (Å) | LIG1$^{EE/AA}$ | LIG1 R771W | LIG1 R641L |
|---|---|---|---|
| LIG1$^{EE/AA}$ K845N | 0.558 (600) | 0.567 (582) | 0.463 (570) |
| LIG1$^{EE/AA}$ | | 0.724 (0.589) | 0.654 (576) |
| LIG1 R771W | | | 0.544 (548) |

**Supplementary Table 1. RMSD values of LIG1 disease mutants.** Table shows RMSD values for the structure of LIG1$^{EE/AA}$ K845N solved in the present study and the structures of LIG1 syndrome mutants that were previously solved by other group (50) for LIG1$^{R641L}$ (7L34) and LIG1$^{R771W}$ (7L35).Values in the parenthesis represents the number of atoms aligned against.

| Nick DNA Substrates | Sequence |
|---|---|
| 3'-dA:A | 5'-CATGGGCGGCATGAACCAGAGGCCCATCCTCACC-3'-FAM<br>3'-GTACCCGCCGTACTTGGACTCCGGGTAGGAGTGG-5' |
| 3'-dG:A | 5'-CATGGGCGGCATGAACCGGAGGCCCATCCTCACC-3'-FAM<br>3'-GTACCCGCCGTACTTGGACTCCGGGTAGGAGTGG-5' |
| 3'-dC:A | 5'-CATGGGCGGCATGAACCCGAGGCCCATCCTCACC-3'-FAM<br>3'-GTACCCGCCGTACTTGGACTCCGGGTAGGAGTGG-5' |
| 3'-dT:A | 5'-CATGGGCGGCATGAACCTGAGGCCCATCCTCACC-3'-FAM<br>3'-GTACCCGCCGTACTTGGACTCCGGGTAGGAGTGG-5' |
| 3'-dT:T | 5'-CATGGGCGGCATGAACCTGAGGCCCATCCTCACC-3'-FAM<br>3'-GTACCCGCCGTACTTGGTCTCCGGGTAGGAGTGG-5' |
| 3'-dG:T | 5'-CATGGGCGGCATGAACCGGAGGCCCATCCTCACC-3'-FAM<br>3'-GTACCCGCCGTACTTGGTCTCCGGGTAGGAGTGG-5' |
| 3'-dC:T | 5'-CATGGGCGGCATGAACCCGAGGCCCATCCTCACC-3'-FAM<br>3'-GTACCCGCCGTACTTGGTCTCCGGGTAGGAGTGG-5' |
| 3'-dA:T | 5'-CATGGGCGGCATGAACCAGAGGCCCATCCTCACC-3'-FAM<br>3'-GTACCCGCCGTACTTGGTCTCCGGGTAGGAGTGG-5' |
| 3'-dA:G | 5'-CATGGGCGGCATGAACCAGAGGCCCATCCTCACC-3'-FAM<br>3'-GTACCCGCCGTACTTGGGCTCCGGGTAGGAGTGG-5' |
| 3'-dT:G | 5'-CATGGGCGGCATGAACCTGAGGCCCATCCTCACC-3'-FAM<br>3'-GTACCCGCCGTACTTGGGCTCCGGGTAGGAGTGG-5' |
| 3'-dG:G | 5'-CATGGGCGGCATGAACCGGAGGCCCATCCTCACC-3'-FAM<br>3'-GTACCCGCCGTACTTGGGCTCCGGGTAGGAGTGG-5' |
| 3'-dC:G | 5'-CATGGGCGGCATGAACCCGAGGCCCATCCTCACC-3'-FAM<br>3'-GTACCCGCCGTACTTGGGCTCCGGGTAGGAGTGG-5' |
| 3'-dA:C | 5'-CATGGGCGGCATGAACCAGAGGCCCATCCTCACC-3'-FAM<br>3'-GTACCCGCCGTACTTGGCCTCCGGGTAGGAGTGG-5' |
| 3'-dT:C | 5'-CATGGGCGGCATGAACCTGAGGCCCATCCTCACC-3'-FAM<br>3'-GTACCCGCCGTACTTGGCCTCCGGGTAGGAGTGG-5' |
| 3'-dC:C | 5'-CATGGGCGGCATGAACCCGAGGCCCATCCTCACC-3'-FAM<br>3'-GTACCCGCCGTACTTGGCCTCCGGGTAGGAGTGG-5' |
| 3'-dG:C | 5'-CATGGGCGGCATGAACCGGAGGCCCATCCTCACC-3'-FAM<br>3'-GTACCCGCCGTACTTGGCCTCCGGGTAGGAGTGG-5' |

**Supplementary Table 2. Nick DNA substrates containing 3'-mismatches.** Nick DNA substrates with 3'-preinserted dA, dT, dG, dC opposite template base A, T, G, or C were used in the ligation assays to investigate the mismatch specificity of LIG1 wild-type and K845N mutant. FAM denotes a fluorescent tag and is located at 3'-end of DNA substrates. The base at the template position is underlined.

| Nick DNA Substrates | Sequence |
|---|---|
| 3'-8oxodG/rG:A | 5'-CATGGGCGGCATGAACC**X**GAGGCCCATCCTCACC-3'-FAM<br>3'-GTACCCGCCGTACTTGGACTCCGGGTAGGAGTGG-5' |
| 3'-8oxodG/rG:C | 5'-CATGGGCGGCATGAACC**X**GAGGCCCATCCTCACC-3'-FAM<br>3'-GTACCCGCCGTACTTGGCCTCCGGGTAGGAGTGG-5' |

**Supplementary Table 3. Nick DNA substrates containing damaged ends.** Nick DNA substrates with 3'-8-oxodG or 3'-8-oxorG opposite template base A or C were used in the ligation assays to investigate the ligation efficiency of LIG1 wild-type and K845N mutant. FAM denotes a fluorescent tag and is located at 3'-end of DNA substrates. The base at the template position is underlined. The damaged base is shown in bold.

| Nick DNA Substrates | Sequence |
|---|---|
| 3'-rA:A | 5'-CATGGGCGGCATGAACC**A**GAGGCCCATCCTCACC-3'-FAM<br>3'-GTACCCGCCGTACTTGGACTCCGGGTAGGAGTGG-5' |
| 3'-rG:A | 5'-CATGGGCGGCATGAACC**G**GAGGCCCATCCTCACC-3'-FAM<br>3'-GTACCCGCCGTACTTGGACTCCGGGTAGGAGTGG-5' |
| 3'-rC:A | 5'-CATGGGCGGCATGAACC**C**GAGGCCCATCCTCACC-3'-FAM<br>3'-GTACCCGCCGTACTTGGACTCCGGGTAGGAGTGG-5' |
| 3'-rG:T | 5'-CATGGGCGGCATGAACC**G**GAGGCCCATCCTCACC-3'-FAM<br>3'-GTACCCGCCGTACTTGGTCTCCGGGTAGGAGTGG-5' |
| 3'-rC:T | 5'-CATGGGCGGCATGAACC**C**GAGGCCCATCCTCACC-3'-FAM<br>3'-GTACCCGCCGTACTTGGTCTCCGGGTAGGAGTGG-5' |
| 3'-rA:T | 5'-CATGGGCGGCATGAACC**A**GAGGCCCATCCTCACC-3'-FAM<br>3'-GTACCCGCCGTACTTGGTCTCCGGGTAGGAGTGG-5' |
| 3'-rA:G | 5'-CATGGGCGGCATGAACC**A**GAGGCCCATCCTCACC-3'-FAM<br>3'-GTACCCGCCGTACTTGGGCTCCGGGTAGGAGTGG-5' |
| 3'-rG:G | 5'-CATGGGCGGCATGAACC**G**GAGGCCCATCCTCACC-3'-FAM<br>3'-GTACCCGCCGTACTTGGGCTCCGGGTAGGAGTGG-5' |
| 3'-rC:G | 5'-CATGGGCGGCATGAACC**C**GAGGCCCATCCTCACC-3'-FAM<br>3'-GTACCCGCCGTACTTGGGCTCCGGGTAGGAGTGG-5' |
| 3'-rA:C | 5'-CATGGGCGGCATGAACC**A**GAGGCCCATCCTCACC-3'-FAM<br>3'-GTACCCGCCGTACTTGGCCTCCGGGTAGGAGTGG-5' |
| 3'-rC:C | 5'-CATGGGCGGCATGAACC**C**GAGGCCCATCCTCACC-3'-FAM<br>3'-GTACCCGCCGTACTTGGCCTCCGGGTAGGAGTGG-5' |
| 3'-rG:C | 5'-CATGGGCGGCATGAACC**G**GAGGCCCATCCTCACC-3'-FAM<br>3'-GTACCCGCCGTACTTGGCCTCCGGGTAGGAGTGG-5' |

**Supplementary Table 4. Nick DNA substrates containing 3'-ribonucleotides.** Nick DNA substrates with 3'-preinserted rA, rG, rC opposite template base A, T, G, or C were used in the ligation assays to investigate the sugar discrimination of LIG1 wild-type and K845N mutant against nick DNA substrates congaining 3'-ribonucleotide. FAM denotes a fluorescent tag and is located at 3'-end of DNA substrates. The base at the template position is underlined. The ribonucleotide at 3'-end of nick is shown in bold.

| Oligonucleotide | Sequence (5'-3') |
| --- | --- |
| Template T | GTCCGACTACGCATCAGC |
| Upstream A | GCTGATGCGT**A** |
| Downstream (5'-P) | P-GTCGGAC |

**Supplementary Table 5. Oligonucleotides used in LIG1 crystallization.** Upstream oligonucleotide (3'-A), downstream oligonucleotide with phosphate (P) at the 5'-end, and template oligonucleotide containing T on a template position were used to prepare the nick DNA substrate with 3'-A:T for LIG1 crystallizations. The base at template base position is underlined and the base position at the 3'-end of nick is shown in bold.

| Oligonucleotide | Sequence |
| --- | --- |
| Up-OH | 5'-Bio-CATGGGCGGCATGAACC**A**-3' |
| Template T | 5'-GGTGAGGATGGGCCTCTGGTTCATGCCGCCCATG-3' |
| Down | 5'-(P)GAGGCCCATCCTCACC-AF488-3' |

**Supplementary Table 6. Nick DNA substrate used in the TIRF.** Up-OH, Template T, and down oligonucleotides were used to prepare the DNA substrate with a single nick site for single-molecule characterization of LIG1 nick DNA binding. Bio denotes a Biotin label located at 5'-end, AF488 is a green-fluorescent dye located at 3'-end, and P stands for a phosphate at 5'-end. The base at 3'-end is shown as bold and the template base is underlined.
